# ISKNV-pORF71 hijacks outer mitochondrial membrane-localized VDAC2 to the nucleus, enhancing apoptosis by disrupting mitochondrial membrane permeability

**DOI:** 10.1101/2025.11.13.688210

**Authors:** He-tong Zhang, Yuting Fu, Youyong Dong, Hemei Qi, Shaoping Weng, Jianguo He, Chuanfu Dong

## Abstract

Infectious spleen and kidney necrosis virus (ISKNV), the type species of the genus Megalocytivirus within the family Iridoviridae, is one of the most devastating pathogens affecting global teleost populations. Our previous study confirmed that ISKNV-pORF71 (p71/VP71) is a viral virulence factor by constructing a recombinant virus with ISKNV orf071 deletion (ISKNV-Δ71). In the present study, we further found that compared with ISKNV-WT, the ISKNV-Δ71 mutant is more capable of maintaining mitochondrial membrane potential, alleviating mitochondrial membrane permeabilization, preserving mitochondrial membrane integrity, sustaining lower cytochrome c release, and thereby reducing the apoptosis rate of infected cells. Further investigations demonstrated a strong interaction between p71 and the outer mitochondrial membrane (OMM)-localized voltage-dependent anion channel 2 (VDAC2). At the subcellular level, p71 and mandarin fish VDAC2 (mfVDAC2) are originally localized to the nucleus and mitochondria, respectively. However, in cells co-transfected with p71 and mfVDAC2, robust nuclear translocation of mfVDAC2 was verified. Endogenous mfVDAC2 was also confirmed to undergo nuclear translocation upon ISKNV-WT infection. In the zebrafish infection model, zebrafish VDAC2 (zfVDAC2)—but not zfVDAC-1 or zfVDAC-3—was observed to translocate into the nucleus through interaction with p71. Using CRISPR/Cas9 technology, VDAC2-knockout zebrafish were successfully constructed. Infection experiments showed that ISKNV-Δ71 exhibits significantly reduced lethality to both wild-type zebrafish and zfVDAC2-knockout zebrafish. In conclusion, our findings reveal that p71 exerts its function during ISKNV infection by hijacking VDAC2 into the nucleus, which alters mitochondrial membrane permeability and enhances ISKNV-induced apoptosis. The phenotypic characteristics of VDAC2 knockout (vdac^⁻/⁻^) zebrafish were also thoroughly analyzed and discussed.

**Author summary:** VDAC2, a well-characterized cellular protein localized to the outer mitochondrial membrane (OMM), plays critical roles in viral pathogenesis and antiviral immune escape. Previous studies have shown that VDAC2 interacts directly or indirectly with viral proteins or functions as a functional receptor to modulate viral pathogenesis; however, few studies have demonstrated that VDAC2 is hijacked from its native OMM to other organelles. Our study here provides robust evidence that cellular VDAC2 is rerouted from its original OMM to the nucleus through interaction with ISKNV p71, thereby altering viral pathogenesis. This work offers novel insights into the role of VDAC2 in pathogen virulence. As far as we know, this is the first report demonstrating that host VDAC2 is hijacked into the nucleus by a viral protein to subsequently modulate the viral life cycle. Collectively, these findings advance our understanding of the active regulation of host functional proteins by iridoviruses.

## Introduction

Members in the family *Iridoviridae* are linear nucleo-cytoplasmic large DNA viruses (NCLDVs) with genome sizes ranging from 98 kb to 212 kp and divided into two subfamilies, namely *Alphairidovirinae* and *Betairidovirinae* (1, 2). The former is mainly composed of three genera including *Lymphocystivirus*, *Ranavirus* and *Megalocytivirus* affecting a variety of cold-blooded vertebrates such as bony fish, amphibians, and reptiles. The latter mainly consists of four genera including *Iridovirus, Chloriridovirus*, *Decapodiridovirus* and *Daphniairidovirus*, infecting invertebrates such as insects, crustaceans, and arthropods, according to the latest updated virus taxonomy of International Committee on Taxonomy of Viruses (ICTV-10) (https://talk.ictvonline.org/taxonomy/). Among *Alphairidovirinae,* members in genus *Megalocytivirus* infect variety of freshwater and marine economically important food fish species and ornamental fish species and has been considered as one of the most dangerous causative agents to bony fish due to its widespread prevalence and high lethality covering numerous countries and regions across a wide geographical range, including Asia, Europe, Australia, North America, and more recently, Africa and South America (3–8). Nowadays, members in genus *Megalocytivirus* are further divided into two species, namely *Megalocytivirus pagrus1* and *M. lates1*, respectively. The former is represented by infectious spleen and kidney necrosis (ISKNV) (9) and the latter is represented by scale drop disease virus (SDDV) (10, 11). The species *M. pagrus1* consists of three genotypes, namely RSIV, ISKNV and TRBIV, and further six sub-genotypes i.e. RSIV-I, RSIV-II, ISKNV-I, ISKNV-II, TRBIV-I and TRBIV-II, respectively (7, 10).

Over the past decade, studies on ISKNV/RSIV have primarily focused on investigating the functions of viral-encoded non-structural viral proteins. For instance, studies have shown that ISKNV VP5 localizes to the inner mitochondrial membrane and exhibits pro-apoptotic activity (12). Viral proteins VP23, VP8 and VP33 interact with Nidogen-1, a key component of the host basement membrane, to form a viral mock-basement membrane. This structure can remodel the extracellular matrix and provide adhesion sites for lymphatic endothelial cells, and it is hypothesized that this mechanism enables the virus to sequester the host immune system for immune evasion (13–15). VP48R is a viral-encoded vascular endothelial growth factor (vVEGF) that stimulates the expression of *flk1*, leading to developmental abnormalities in zebrafish (*Danio rerio*) embryos, including pericardial edema and tail expansion (16). VP111L interacts with TNFR-associated death domain protein (TRADD) and induces cellular apoptosis (17). VP119L binds to the host PINCH protein to inhibit integrin-linked kinase (ILK) signaling and interfere with embryonic angiogenesis in zebrafish (18).

To date, 38 and 44 viral structural proteins and have been identified in ISKNV and RSIV virions, respectively, but the roles of these structural proteins in viral replication or virus-host interactions have rarely been clarified (19, 20). Our recent studies have shown that the knockout mutant (ISKNV-Δ71) of the ISKNV structural protein VP71, generated via recombinant virus technology, exhibits higher replication efficiency but significantly reduced lethality compared to the parental strain (21). Partial data indicate that VP71 may be involved in ISKNV infection-induced apoptosis, though the underlying mechanism remains unclear. On the other hand, our previous viral proteomics studies have also identified several important host proteins—such as heat shock proteins 90 and 70 (HSP90, HSP70), virus-inducible stress protein (VISP), voltage-dependent anion channel 2 (VDAC2), and valosin containing protein (VCP/p97)—that are directly involved in the packaging of mature virions (20). This finding suggests that these host proteins may either actively participate in or be manipulated by the virus to contribute to the viral life cycle, although their specific functional mechanisms remain poorly studied.

VDAC, also known as mitochondrial porins, are the most abundant proteins in the outer mitochondrial membrane (22). In mammals, three VDAC isoforms have been identified, sharing approximately 75% nucleotide sequence similarity and a molecular weight of 32–34 kDa. Despite high nucleotide and amino acid sequence similarity among all three VDAC isoforms, they exhibit certain functional redundancy (23). VDAC1 is expressed at the highest level among the three isoforms in most cell types (24), and functions in the transport of metabolites such as ATP/ADP, NAD/NADH, Ca²⁺, and other small molecules. In addition to its transport function, VDAC2 plays a role in regulating apoptosis (25). VDAC3 exhibits weaker membrane insertion and pore-forming activity, suggesting that its primary function is not as an ion channel (26). Instead, it has been found to localize to centrioles and regulate centriole assembly (27), and is also demonstrated to be present in the outer dense fibers of sperm, participating in maintaining sperm motility (28).

Cellular apoptosis is triggered by three distinct signaling pathways, namely the extrinsic pathway, intrinsic pathway, and Granzyme B (GrB) pathway. All three pathways can transmit apoptotic signals to mitochondria, initiating apoptosis through mitochondrial membrane permeabilization (MMP). Currently, two major mechanisms are considered to induce MMP: (a) Mitochondrial Outer Membrane Permeability (MOMP) occurs at the outer mitochondrial membrane (OMM). Certain Bcl-2 family proteins (e.g., Bax and Bak) assemble into large homomeric or heteromeric multimeric channels, which mediate the release of mitochondrial intermembrane space proteins to activate apoptosis (29–31). During MOMP, the mitochondrial inner membrane typically remains intact and sustains respiratory metabolic activity for an extended period, allowing mitochondria to retain functionality and produce ATP (32). (b) Mitochondrial Permeability Transition (MPT) occurs at the inner mitochondrial membrane (IMM) and is mediated by the mitochondrial permeability transition pore (MPTP). This large pore complex spans both the inner and outer mitochondrial membranes, permitting the passage of substances with a molecular weight < 1.5 kDa. Prolonged opening of MPTP leads to the loss of mitochondrial membrane potential, cessation of ATP synthesis, and mitochondrial swelling and rupture (33). The composition of MPTP remains controversial to date. Literatures from the late 1990s to the early 2000s proposed that MPTP is formed through complex interactions between Bcl-2 family members, the outer mitochondrial membrane protein VDAC, the inner membrane protein adenine nucleotide translocators (ANT), and cyclophilin D (CypD) located in the mitochondrial matrix (34, 35). However, subsequent studies demonstrated that mitochondria from cells with knockout of all three VDAC isoforms retain MMP capability. Additionally, *vdac2* knockout in cells does not inhibit MMP but instead exacerbates cell death (25, 36), indicating that VDAC is not an essential component of MPTP but plays a regulatory role in its activity.

Our previous study confirmed that both ISKNV-p71 and VDAC2, as structural proteins, are involved in the formation of mature virions (20). In the present study, we further demonstrated that p71 can manipulate VDAC2 to translocate into the nucleus, thereby affecting viral pathogenicity. This is the first report on the critical biological event where VDAC2 is hijacked into the nucleus by a virus, which provides new insights into a comprehensive understanding of the biological functions of VDAC2.

## Materials and methods

### Animal ethics statement

All animal experiments were performed in accordance with the regulations for animal experiments of Guangdong Province, China, and were permitted by the Ethics Committee of Sun Yat-sen University with protocol codes: SYSU-IACUC-2018-000057 and SYSU-IACUC-2021-000326.

### Viruses, cell lines, and antibodies

The mandarin fish fry (MFF-1) cell line was cultured in high-glucose DMEM medium supplemented with 10% fetal bovine serum (FBS) in an incubator at 27°C with 5% CO₂ (37). BHK21 and HeLa cells were obtained from ATCC and cultured in DMEM medium containing 10% FBS at 37°C in a 5% CO₂ incubator. The wild-type ISKNV-NH060831 (ISKNV-WT) was isolated and characterized from diseased mandarin fish in 2006 and preserved in our laboratory (19, 37). The ISKNV *orf71* deletion mutant (ISKNV-Δ71) was previously constructed and stored in our laboratory (21). MFF-1 cells were infected with the virus at a multiplicity of infection (MOI) of 1. Cell cultures were collected at 5 days post infection (dpi), and viral particles were released by three freeze-thaw cycles. The virus stock was stored at -80°C until use.

Polyclonal or monoclonal antibodies against antiviral proteins (p6, p7, p23, p62, p71, p101) have been described in our previous reports (8, 19). Tag-specific antibodies (anti-HA, -GFP, -Flag, -Myc, etc.) were purchased from a commercial antibody company (ABMART, Shanghai).

### Flow cytometric analysis of apoptosis

The operation procedure of the BD FACSMelody flow cytometer was performed according to the manufacturer’s instructions. For cells overexpressing exogenous proteins: Apoptotic cells were labeled using the Annexin V-FITC Apoptosis Detection Kit (Sigma-Aldrich). Labeled cells were detected using the FL1 channel (BD C6 flow cytometer) or FITC channel (BD FACSMelody flow cytometer). Detection of apoptosis induced by ISKNV-WT or ISKNV-Δ71: MFF-1 cells were infected with ISKNV-WT or ISKNV-Δ71 at an MOI of 0.1, respectively. Apoptotic cells were labeled with Annexin V Alexa Fluor 555 conjugate and detected using the PE channel (BD FACSMelody flow cytometer). Positive control for apoptosis induction: Apoptosis was induced with staurosporine at a final concentration of 1 μg/mL, and detection was performed 1 hour after drug addition.

### Prokaryotic expression of *mf*VDAC2 and antibody preparation

The *mfvdac2* gene sequence was obtained by RACE-PCR clone (S1 Fig) and was constructed recombinant plasmids with pMAL expression system. Briefly, the recombinant plasmids were transformed into BL21 competent cells, and the transformation method was performed as product instruction. Single colonies of the expression strain were picked and inoculated into 20 mL of LB medium containing the appropriate antibiotic, followed by shaking incubation at 37℃ overnight. On the next day, the bacterial culture was sub-cultured into fresh LB medium at a 1:100 ratio. When the optical density at 600 nm (OD₆₀₀) of the bacterial suspension reached 0.6-0.8, isopropyl β-D-1-thiogalactopyranoside (IPTG) was added at a final concentration of 1 mM (1:1,000 dilution ratio), and induction was carried out for 4-6 hours. A 1 mL aliquot of the bacterial culture was harvested by centrifugation at 5,000×g and 4℃ for 10 minutes. The bacterial pellet was resuspended in 50 μL of PBS, and 10 μL of 5× SDS-PAGE loading buffer was added. After mixing thoroughly, the sample was incubated in a 95℃-water bath for 10 minutes. Following SDS-PAGE electrophoresis, the gel was stained with Coomassie Brilliant Blue R-250 for 30 minutes and destained overnight to analyze the protein expression level. Sufficient purified fusion protein was subjected to SDS-PAGE, and the target protein band was excised from the gel. The gel slice was ground thoroughly and mixed with Complete Freund’s Adjuvant (CFA) for full emulsification, which was used as the antigen for the primary immunization of Balb/c mice. The immunization dose was 80-150 μg per mouse, administered via subcutaneous multi-point injection. The second immunization was performed 14 days later, using the same antigen dose emulsified with Incomplete Freund’s Adjuvant (IFA). The third to fifth immunizations were conducted at 10-day intervals, with the same antigen dose and Incomplete Freund’s Adjuvant used for each. Three days after the fifth immunization, blood was collected from the mice by enucleation into 1.5 mL centrifuge tubes. The blood samples were allowed to stand at 37℃ for 1 hour and then at 4℃ overnight. On the following day, the serum was collected by centrifugation at 15,000×g and 4℃ for 15 minutes and stored at -80℃ until use.

### SDS-PAGE and Western-blotting (WB) assay

SDS-PAGE and WB assays were performed as previously described by our group (19, 37, 38). The purified ISKNV virion was referred to our previous description (19, 20). For fractionation of virion proteins, the purified ISKNV was treated by 1% Triton X-100 for 3 min at room temperature, and were separated by centrifugation at 20,000 g for 30 min at 4°C. The supernatant and pellet were collected for SDS-PAGE and WB analysis as previous report (19).

### Measurement of mitochondrial membrane potential was using the JC-1 probe

JC-1 (Thermo Scientific) was dissolved in dimethyl sulfoxide (DMSO) to prepare a 2 mg/mL stock solution, which was stored at -20 ℃. Prior to laser confocal imaging, the JC-1 probe was added to a final concentration of 2.0 µg/mL, followed by incubation at 27℃ for 30 min. Cells were washed three times with sterile phosphate-buffered saline (PBS), and confocal imaging was performed immediately. Mitochondrial membrane permeability was measured using Calcein AM. The stock solution of Calcein AM (Thermo Scientific) was diluted with PBS to a final concentration of 2 µM. The culture medium was replaced with this diluted Calcein AM solution, and the cells were incubated at 27℃ for 30 min. After three washes with PBS, cobalt chloride (CoCl₂) was added to a final concentration of 400 µM. The cells were incubated for 15 min and then imaged under a confocal microscope.

### Effect of DIDS on ISKNV replication

4,4′-Diisothiocyanatostilbene-2,2′-disulfonic acid disodium salt hydrate (DIDS; Sigma-Aldrich) was dissolved in DMSO to prepare a 0.01 M stock solution. For use, the stock solution was diluted with culture medium to a final concentration of 100 µM. One hour after ISKNV infection, the original medium was replaced with DIDS-containing medium, and the cells were transferred to an incubator for further culture.

### Mitochondrial isolation

Mitochondria were isolated using the Mitochondria Isolation Kit for Cultured Cells (Thermo Scientific, USA) following the manufacturer’s protocol. Cells cultured in 100 mm² dishes were harvested by trypsinization and collected via centrifugation at 850×g for 2 minutes, yielding approximately 2×10⁷ cells. Subsequently, 800 µL of Mitochondrial Isolation Reagent A was added to the cell pellet, followed by vortexing at medium speed for 5 seconds and incubation on ice for 2 minutes. Next, 10 µL of Mitochondrial Isolation Reagent B was added, and the mixture was vortexed at maximum speed for 5 seconds. The centrifuge tube was then incubated on ice for 5 minutes, with vortexing at maximum speed for 5 seconds every minute during incubation. Following this, 800 µL of Mitochondrial Isolation Reagent C was added, and the sample was mixed by inversion (avoid vortexing). The mixture was centrifuged at 700×g and 4°C for 10 minutes. The supernatant was transferred to a new centrifuge tube and centrifuged at 3000×g and 4°C for 15 minutes. The resulting supernatant (cytosolic fraction) was transferred to a fresh tube, while the pellet contained the isolated mitochondria. To further purify the mitochondria, 500 µL of Mitochondrial Isolation Reagent C was added to the pellet, followed by centrifugation at 12,000×g and 4°C for 5 minutes. The supernatant was discarded, and the final pellet consisted of purified mitochondria.

### Transmission electron microscopy (TEM)

MFF-1cells infected with ISKNV-WT or ISKNV-Δ71 was collected and fixed with 2.5% glutaraldehyde in 0.1M PBS (pH 7.4) and post-fixed in 0.1M PBS containing 2.0% osmium tetroxide. Ultrathin sections were stained with uranyl acetate-lead citrate and examined by a Philips CM10 electron microscopy as our previous description (4, 38).

### Co-Immunoprecipitation (Co-IP) assay

Forty-eight hours post-transfection, HeLa cells were washed three times with pre-chilled PBS. Merck protease inhibitor cocktail was added to the IP lysis buffer at a 1:100 dilution ratio. The lysis buffer was then added to the cell culture dishes, and cells were scraped off on ice using a cell scraper. The cell suspension was transferred to 1.5 mL centrifuge tubes and lysed on ice for 30 minutes. Following lysis, the samples were centrifuged at 15,000×g and 4℃ for 10 minutes, and the supernatant was transferred to fresh 1.5 mL centrifuge tubes. Thermo Dynabeads were prepared by vortexing for 15 seconds to resuspend the beads thoroughly. A 50 μL aliquot of the beads was pipetted into a 1.5 mL centrifuge tube, which was then placed on a magnetic rack to discard the supernatant. The beads were washed once with PBS containing Tween-20 (PBST). A predetermined amount of primary antibody was diluted in 200 μL of PBST and mixed with the beads, followed by incubation on a rotating mixer at room temperature for 30 minutes. The tube was placed on the magnetic rack to remove the supernatant, and the beads were washed three times with 500 μL of PBST (5 minutes each wash). Subsequently, 300 μL of the cell lysate (obtained from the aforementioned centrifugation step) was added to the bead-antibody complex, and the mixture was incubated with rotation at room temperature for 1 hour. After incubation, the tube was placed on the magnetic rack to discard the supernatant, and the beads were washed three times with 500 μL of PBST (5 minutes each wash). Finally, 50 μL of 1× SDS-PAGE loading buffer was added to the beads, and the mixture was vortexed thoroughly before being incubated in a 95 ℃ water bath for 10 minutes. The samples were then subjected to WB analysis.

### Subcellular localization analysis of fluorescent fusion proteins

MFF-1, BHK21 or HeLa cells were seeded in 35 mm glass-bottom dishes and transfected upon reaching 80% confluency. Thirty-six hours post-transfection, cells were washed three times with pre-chilled PBS. Hoechst 33342 nuclear stain (diluted at 1:2000) was added, and the cells were incubated on ice for 5 minutes. The stain was then replaced with pre-chilled PBS, and the cells were observed under a laser confocal microscope.

### Indirect immunofluorescence assay (IFA)

BHK21 or HeLa cells were seeded in 35 mm glass-bottom dishes and transfected when reaching 80% confluency. Thirty-six hours post-transfection, cells were washed three times with pre-chilled phosphate-buffered saline (PBS). Pre-chilled paraformaldehyde (PFA) was added to fix the cells on ice for 10 minutes, followed by three washes with PBS (5 minutes each). Subsequently, 500 μL of 1% Triton X-100 was added for permeabilization at room temperature for 10 minutes, and cells were rinsed three times with PBS (5 minutes each). For blocking, 200 μL of 10% normal goat serum (NGS) at a final concentration was added, and the cells were incubated at 37℃ for 1 hour. Next, rabbit anti-HA primary antibody and mouse anti-V5 primary antibody (both diluted at 1:500) were added, and the cells were incubated overnight at 4℃. On the following day, cells were washed three times with PBS (5 minutes each). Goat anti-rabbit IgG conjugated to Alexa Fluor 488 and donkey anti-mouse IgG conjugated to Alexa Fluor 555 (secondary antibodies) were then added, and the cells were incubated at 37℃ for 1 hour, followed by three washes with PBS (5 minutes each). Hoechst 33342 nuclear stain (diluted at 1:2000) was added, and the cells were incubated at room temperature for 10 minutes. After three final washes with PBS (5 minutes each), the cells were analyzed using a laser confocal microscope.

### Yeast two hybrid (Y2H)

*mf*VDAC2-BD was used as the bait protein, and 23 ISKNV viral proteins served as target proteins. The transformation of Y2H plasmids was performed with reference to the instructions of the Yeast maker Yeast Transformation System 2 kit, and the specific procedures are briefly described as follows: The constructed recombinant plasmids (VDAC2-pGBKT7 bait plasmid and ISKNV-encoded protein-pGADT7 target plasmid) were thoroughly mixed with 50 μg of yeast carrier DNA, and then added to competent yeast cells. The amount of recombinant plasmid in each tube was 100 ng. Meanwhile, positive and negative controls were set up to verify the validity of the experiment. The carrier DNA was first heated at 100℃ for 5 min, then quickly cooled in an ice bath for denaturation, and this denaturation process was repeated once. 500 μL of PEG/LiAc solution was added to the system, mixed gently, and then incubated at 30℃ for 30 min. During the incubation, the centrifuge tube was inverted once every 10 min to keep the system mixed. 20 μL of DMSO was added, mixed gently, and then heat-shocked in a 42℃ -water bath for 15 min. The centrifuge tube was inverted once every 5 min to ensure uniform heat shock. After centrifugation at 12000 r/min for 15 s, the supernatant was discarded. The bacterial pellet was resuspended in 1 mL of LYPD Plus Medium and incubated at 30℃ for 90 min to restore bacterial viability. Centrifugation was performed again at 12000 r/min for 15 s, the supernatant was discarded, and the bacteria were resuspended in 1 mL of 0.9% (w/v) NaCl solution. The resuspended bacterial solution was diluted 10-fold, and 100 μL of the diluted bacterial solution was evenly spread on the SD/-Leu/-Trp double-deficient medium plate, followed by constant temperature culture at 30℃. After approximately 5 days of culture, single colonies appeared on the plate. Red single colonies with a diameter greater than 2 mm were selected and transferred to the SD/-Leu/-Trp/-His triple-deficient medium plate, and constant temperature culture was continued at 30℃. When clones appeared on the triple-deficient plate, single clones were picked and spread on the SD/-Leu/-Trp/-His/-Ade/X-α-gal quadruple-deficient chromogenic medium plate, followed by constant temperature culture at 30℃. After 5 days, the growth of blue bacteria on the plate was observed to determine the interaction between proteins.

### Zebrafish source, rearing and breeding

Zebrafish broodstock were purchased from the China Zebrafish Resource Center (CZRC). The zebrafish were reared in a constant-temperature circulating aquatic system at 28℃, which was equipped with physical filtration units and ultraviolet sterilization units. A light-dark cycle of 14 h light and 10 h dark was adopted. One day before breeding, female and male fish were placed in a spawning tank at a ratio of 1:1 or 1:2, and a partition was used to separate females from males. On the next morning at 8:00, light was provided, the partition was removed, and the zebrafish began to mate. The collected fish eggs were washed twice with rearing water. Embryos were incubated in rearing water containing 0.5 mg/L methylene blue at 28 ℃. Larvae hatched at approximately 4 days post-fertilization (dpf), and no feeding was required during this stage. Zebrafish were at the larval stage from 5 to 15 dpf, during which they were fed with paramecium and powdered feed. After 15 dpf, they could be fed with brine shrimp, juvenile powdered feed and other baits.

### Gene editing (gRNA design, vector construction and embryo injection)

The gRNA was designed via an online website (http://zifit.partners.org/) to minimize off-target effects, with the gRNA targeting sequence of GTGGAATACCGACAACACCC. The designed gRNA sequence was cloned into the pDR274 plasmid (Addgene Plasmid #42250) through the BsaI restriction enzyme site, and in vitro transcription was performed together with pCS2-nCas9n (Addgene Plasmid #42251) using the mMACHINE T7 and mMACHINE SP6 kit (Invitrogen). Each embryo was injected with 20 μL of mixed RNA, containing 25 ng/μL gRNA and 200 ng/μL Cas9 mRNA. Zebrafish were anesthetized with tricaine, and a caudal fin tissue sample of approximately 2 mm² was cut off using a scalpel. DNA was extracted using the QIAGEN DNeasy Blood & Tissue Kit, followed by PCR detection. Sequencing peak maps were analyzed to identify the type of gene editing.

### Virus challenge, histopathology and immunofluorescence

Zebrafish were anesthetized with tricaine and intraperitoneally injected using a microsyringe (100 μL capacity). The injection dose was 10¹⁰ copies/ml, with 20 μL per fish. ISKNV-WT was injected into wild-type zebrafish and *vdac2^⁻/⁻^* zebrafish, respectively; ISKNV-Δ71 was injected into wild-type zebrafish and *vdac2^⁻/^*^⁻^ zebrafish, respectively. For each virus challenge group, the injection dose was 10¹⁰ copies/ml (20 μL per fish via intraperitoneal injection). For the four challenge combinations, liver and spleen tissues were sampled at 4 dpi for tissue sectioning and immunofluorescence analysis. The mock group was injected with MFF-1 cell culture. Primary antibodies used were mouse monoclonal antibody against ISKNV-VP23 (8) and rabbit polyclonal antibody against ISKNV MCP (19). Secondary antibodies used were goat anti-mouse antibody conjugated with Alexa Fluor 488 and goat anti-rabbit antibody conjugated with Alexa Fluor 555. Images were captured using a Leica SP8 confocal laser scanning microscope.

### Primer sequence data

The details of the relevant primer sequences involved in experiments such as molecular cloning and expression, quantitative PCR (qPCR) analysis, yeast two-hybrid (Y2H) assay, and construction of various recombinant plasmids are provided in Supplementary Table 1 (S1 Table).

### Statistical analysis

All data are presented as the mean ± standard deviation (SD) from three independent experiments.

Statistical analyses were performed using SPSS 16.0 software. Student’s t-test or one-way analysis of variance (ANOVA) was employed for comparisons between the control group and experimental groups, depending on the number of groups involved. Statistical significance was defined as follows: *P < 0.05 (significant), **P < 0.01 (highly significant), and differences with P ≥ 0.05 were considered nonsignificant and marked as “ns”. Additionally, different lowercase letters (a, b, c, d, etc.) were used to label groups, where distinct letters indicate significant differences between groups.

## Results

### ISKNV-WT induces more robust apoptosis in MFF-1 cells than ISKNV-Δ71

It has been reported that apoptosis induced by ISKNV/RSIV infection may contribute to the cytopathic effect (CPE) (37, 39, 40). Given that ISKNV-Δ71 infection accelerates CPE in MFF-1 cells (21), we investigated whether VP71 knockout alters cell apoptosis. We first purified ISKNV-WT and ISKNV-Δ71. SDS-PAGE analysis showed that both viruses were well-purified (**Fig. 1A**). WB analysis demonstrated that the p71 protein was present in purified ISKNV-WT but undetectable in purified ISKNV-Δ71, confirming that p71 was indeed knocked out in ISKNV-Δ71 (**Fig. 1A**). Then, MFF-1 cells were infected with ISKNV-WT or ISKNV-Δ71 at an MOI of 0.1, and apoptosis was detected at 24 hours post-infection (hpi), 48 hpi, and 72 hpi. Apoptotic cells were labeled with Annexin V conjugated to Alexa Fluor™ 555 to avoid interference from EGFP expressed by ISKNV-Δ71, followed by flow cytometric analysis. The results showed that both viruses induced apoptosis, with the apoptosis rate increasing over time. At 24 hpi, the apoptosis rates induced by the two viruses were comparable: 18.38% for ISKNV-WT and 15.2% for ISKNV-Δ71. However, a significant difference in apoptosis rate emerged at 48 hpi (33.81% for ISKNV-WT vs. 25.45% for ISKNV-Δ71) and became more pronounced at 72 hpi (48.72% for ISKNV-WT vs. 40.28% for ISKNV-Δ71). These findings demonstrate that ISKNV-Δ71 induces a lower apoptosis rate in infected cells compared to ISKNV-WT (48 hpi: P=0.0137; 72 hpi: P=0.0029) (**Fig. 1-B and C**).

**Figure 1.**
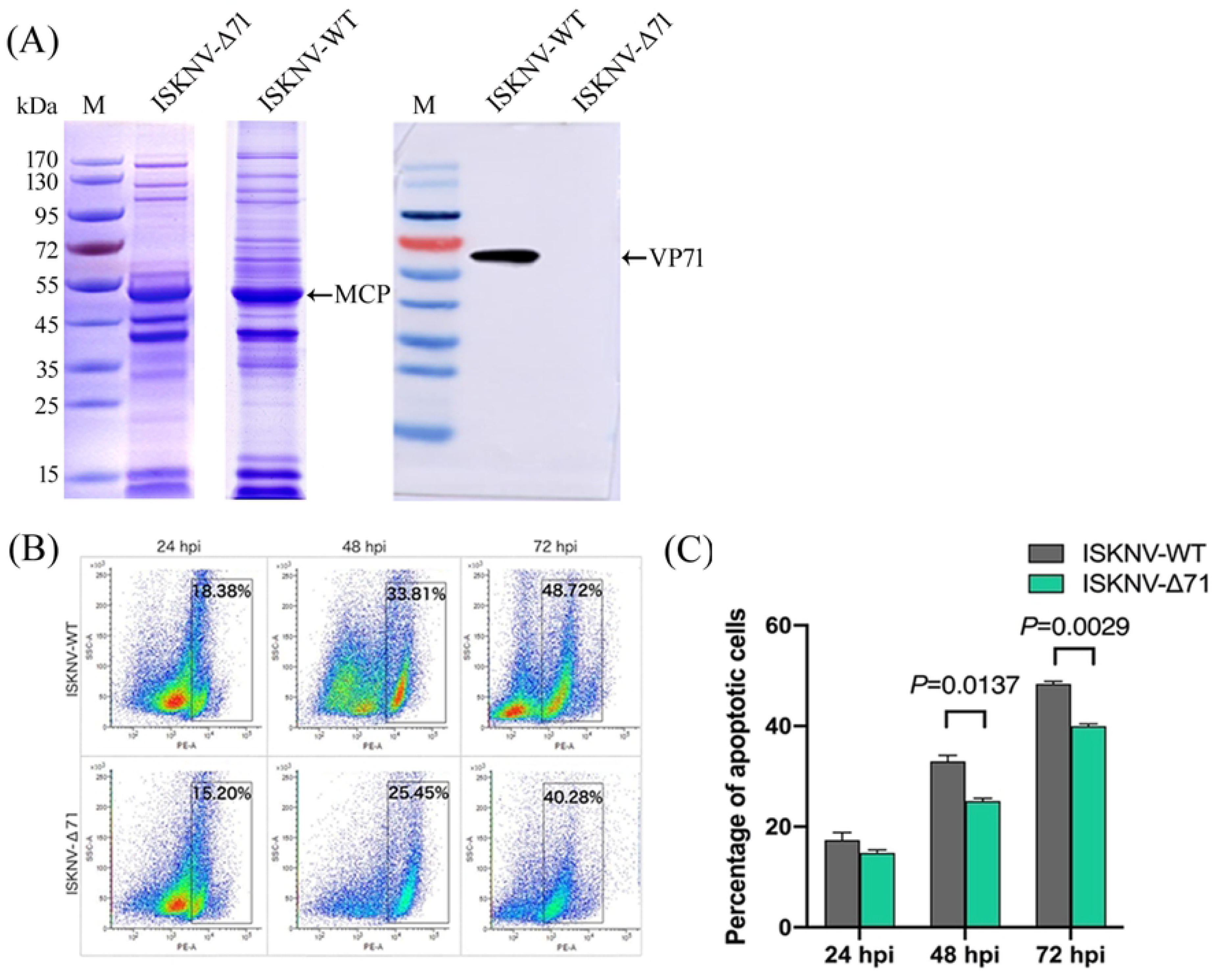
The *orf71*-knockout ISKNV (ISKNV-Δ71) significantly reduces the apoptosis of infected cells compared with the parental ISKNV strain (ISKNV-WT). (A), SDS-PAGE and WB analyses were performed to examine the structural proteins of purified ISKNV-Δ71 and its parental ISKNV-WT; (B), Flow cytometry was used to compare and analyze the apoptosis of MFF-1 cells induced by ISKNV-Δ71 and ISKNV-WT infection; C. Statistical analysis showed that ISKNV-Δ71 significantly decreased the cell apoptosis rate compared with that of ISKNV-WT.

### ISKNV-WT significantly alters the mitochondrial membrane potential (ΔΨm) of infected cells compared to ISKNV-Δ71

We characterized the ΔΨm using JC-1 (41). JC-1 is a cationic carbocyanine dye that accumulates more in hyperpolarized mitochondria (hyperpolarization indicates the inner mitochondrial membrane has a transmembrane potential, with the mitochondrial matrix being electronegative) to form aggregates, emitting red fluorescence (590 nm). Depolarized mitochondria (depolarization indicates a decrease or loss of inner mitochondrial membrane potential) accumulate less JC-1, which then exists as monomers emitting green fluorescence (535 nm). MFF-1 cells were infected with ISKNV at an MOI of 0.1. At 48 hpi, the original medium was replaced with medium containing the JC-1 probe. After incubation for 30 minutes, cells were observed under a confocal fluorescence microscope. In the control group, mitochondria mainly appeared linear or short rod-shaped—their normal morphology—with distinct red fluorescence representing high transmembrane potential. Bright-field images showed cells adhered well with normal morphology (**Fig. 2-g, h, i**). In ISKNV-WT-infected cells, red fluorescence was reduced, indicating decreased mitochondrial transmembrane potential. Mitochondria appeared as discrete green bright spots, which may suggest mitochondrial swelling (**Fig. 2-a, b, c**). In contrast, mitochondria in ISKNV-Δ71-infected cells appeared as red bright spots with clear boundaries, demonstrating that mitochondrial transmembrane potential was maintained (**Fig. 2-d, e, f**). The preserved ΔΨm in ISKNV-Δ71-infected MFF-1 cells indicates the integrity of the inner mitochondrial membrane. The decreased ΔΨm in ISKNV-WT-infected cells may result from mitochondrial dysfunction (42, 43), opening of mitochondrial permeability transition pores, loss of electrochemical gradient, or other factors. The underlying cause of this change may be the reduced levels of VDAC2, a protein localized to the outer mitochondrial membrane (44).

**Figure 2.**
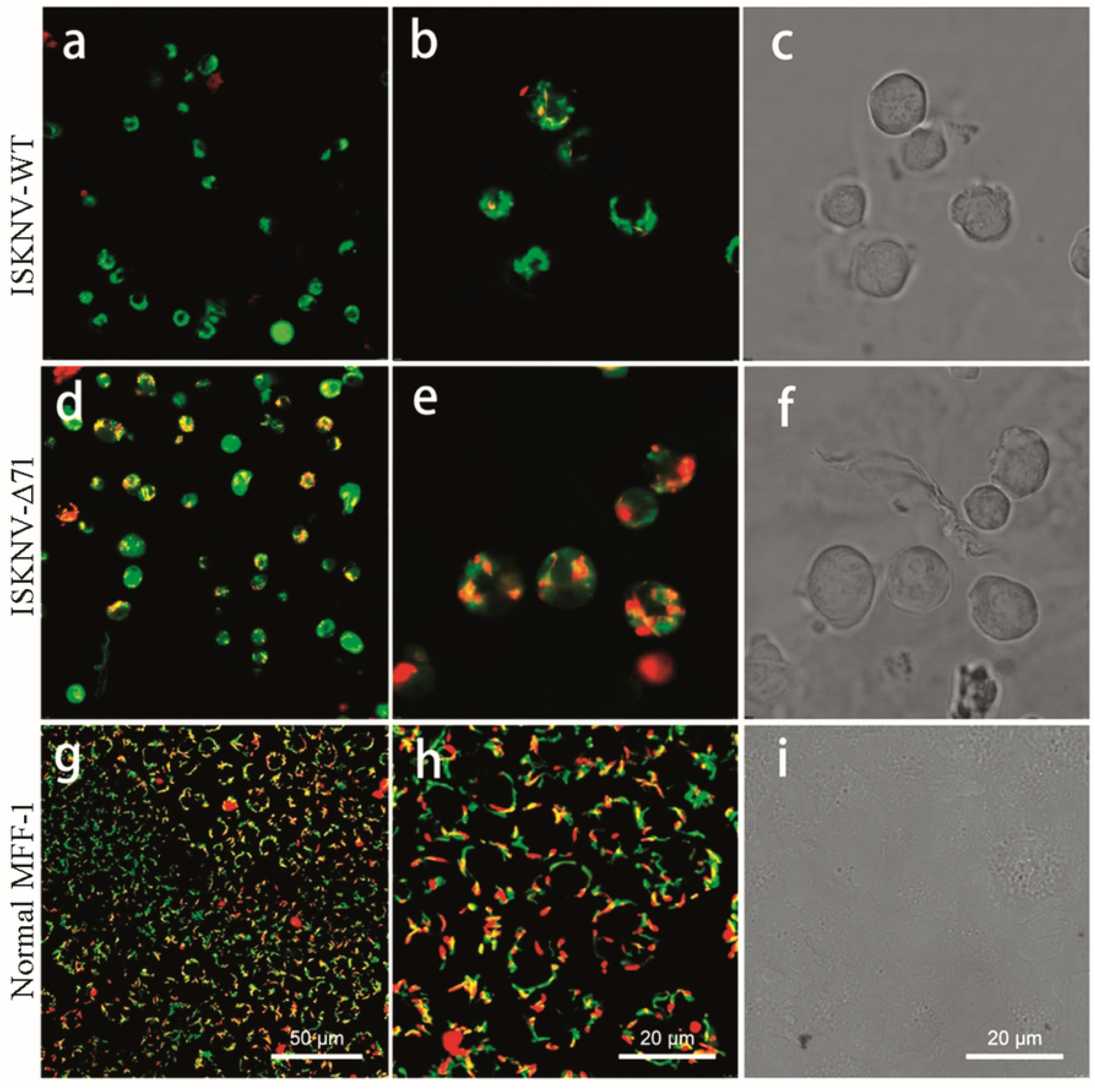
The mitochondrial membrane potential (ΔΨm) in MFF-1 cells infected with ISKNV-WT is lower than that in MFF-1 cells infected with ISKNV-△71. JC-1 probe was used to determine the mitochondrial inner membrane potential of MFF-1 cells infected with ISKNV-WT and ISKNV-Δ71 at an MOI of 0.1 at 48 hpi. The probe emits red fluorescence when **ΔΨm** is high, and green fluorescence when the potential is low. The scale bar is 50 μm or 20 μm. (a) Fluorescence image of ISKNV-WT infection (40×). (b, c) Fluorescence image and corresponding bright-field image of ISKNV-WT infection (100×). (d) Fluorescence image of ISKNV-Δ71 infection (40×). (e, f) Fluorescence image and corresponding bright-field image of ISKNV-Δ71 infection (100×). (g) Fluorescence image of the control group (40×). (h, i) Fluorescence image and corresponding bright-field image of the control group (100×).

### ISKNV-WT is more capable of altering mitochondrial membrane permeability by disrupting mitochondrial membrane integrity compared to ISKNV-Δ71

We used calcein acetoxymethyl ester (Calcein AM) to detect differences in mitochondrial permeability between ISKNV-WT and ISKNV-Δ71 infections. Calcein AM can passively diffuse into cells and accumulate in the cytoplasm, including mitochondria (45, 46). In ISKNV-Δ71-infected MFF-1 cells exhibiting CPE, more fluorescence was retained compared to ISKNV-WT-infected cells (**Fig. 3A**), with cell outlines marked by white dashed lines. No fluorescence loss was observed in the control group. These results indicate that ISKNV-WT infection increases mitochondrial membrane permeability in MFF- 1 cells, although a small number of mitochondria retain membrane integrity; in contrast, more mitochondria maintain membrane integrity in ISKNV-Δ71-infected MFF-1 cells. Notably, both ISKNV strains induced varying degrees of increased mitochondrial membrane permeability compared to the control group (**Fig. 3A**). Increased mitochondrial membrane permeability ultimately leads to mitochondrial swelling and rupture. Therefore, we infected MFF-1 cells with ISKNV-WT or ISKNV-Δ71 and observed mitochondria using TEM. At 24 and 48 hpi, some mitochondria in ISKNV-WT-infected cells appeared swollen, with a small number undergoing rupture—a phenomenon consistent with observations in other iridoviruses (47). In contrast, mitochondria in ISKNV-Δ71-infected cells maintained a clear, intact elliptical morphology. By 120 hpi, a large number of mitochondria in ISKNV-WT-infected MFF-1 cells were ruptured, with mitochondrial cristae barely distinguishable. In ISKNV-Δ71-infected cells, only a small number of mitochondria were ruptured, and some retained a high-electron-density matrix and indistinct cristae (**Fig. 3B**). Thirty random TEM fields (2000× magnification) were observed to count ruptured mitochondria (samples from 72 hpi and 120 hpi, with 15 fields observed per strain per time point). The results showed that 66.8% of mitochondria in ISKNV-WT-infected MFF-1 cells were ruptured, compared to 39.7% in ISKNV-Δ71-infected cells (**Fig. 3C**). This demonstrates that mitochondrial integrity is stronger in ISKNV-Δ71-infected cells.

**Figure 3.**
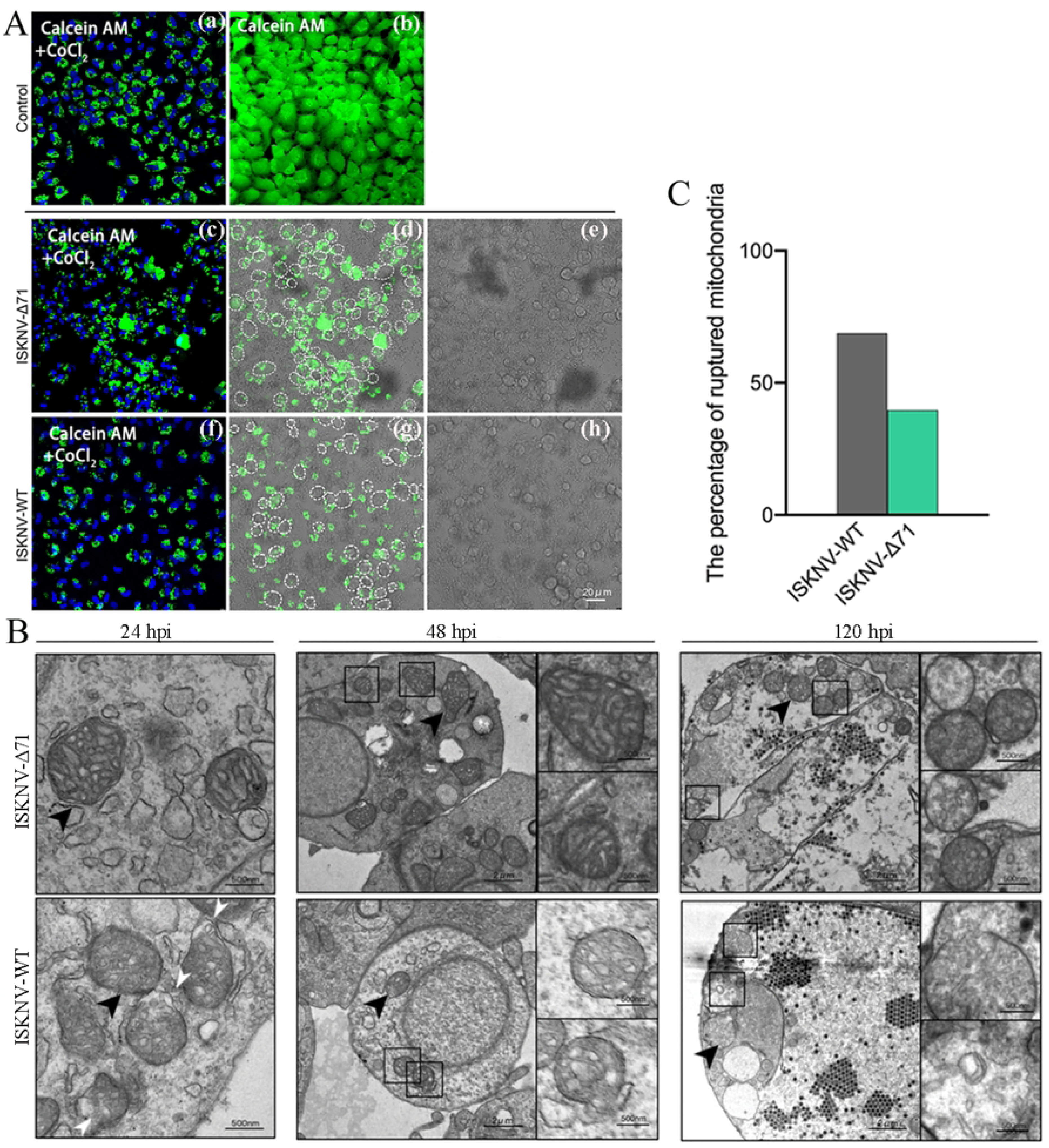
The mitochondrial membrane permeability in MFF-1 cells infected with ISKNV-WT is higher than that in MFF-1 cells infected with ISKNV-△71. (A), Mitochondria with intact membranes exhibit green fluorescence emitted by Calcein, while the fluorescence in mitochondria with membrane permeabilization is reduced or absent. Compared with the control group (a and b), partial mitochondria in ISKNV-△71-infected cells show membrane permeabilization (c-e); by contrast, ISKNV-WT infection leads to more severe mitochondrial membrane permeabilization (f-h). (B), TEM-based comparative analysis of mitochondrial integrity in MFF-1 cells infected by ISKNV-WT and ISKNV-Δ71. Partial mitochondria in ISKNV-WT-infected cells were swollen and ruptured (white arrowheads) at 24 and 48 hpi, with extensive mitochondrial rupture observed at 120 hpi, where mitochondrial cristae were barely distinguishable. In ISKNV-Δ71-infected cells, mitochondria maintained normal morphology at 24 and 48 hpi. Even at 120 hpi, electron-dense matrix remained detectable, and mitochondrial cristae were faintly visible. (C), Combined statistical analysis of mitochondrial rupture at 72 and 120 hpi revealed that the mitochondrial rupture rate of ISKNV-WT was 27.1% higher than that of ISKNV-Δ71

### ISKNV-WT facilitates the release of mitochondrial cytochrome c more effectively than ISKNV-Δ71

Cytochrome c is an intermembrane space protein of mitochondria; outer mitochondrial membrane permeabilization leads to its release into the cytosol. Subsequently, cytochrome c mediates the allosteric activation of apoptotic protease activating factor-1 (APAF-1), which induces the proteolytic maturation of caspase-9 and activates the caspase cascade. Activated caspases ultimately trigger cell apoptosis (48). We detected cytochrome c release following mitochondrial integrity disruption. MFF-1 cells were infected with ISKNV at an MOI of 0.1, and samples were collected at different time points for WB analysis. The level of cytochrome c released from mitochondria to the cytoplasm increased with infection time in ISKNV-WT-infected cells, reaching 2.5-fold that of the control group at 3 dpi. Cytochrome c release also showed an upward trend in ISKNV-Δ71-infected cells, but the increase was less pronounced compared to ISKNV-WT, with the peak value (1.5-fold that of the control group) observed at 5 dpi (**Fig. 4A**).

**Fig. 4.**
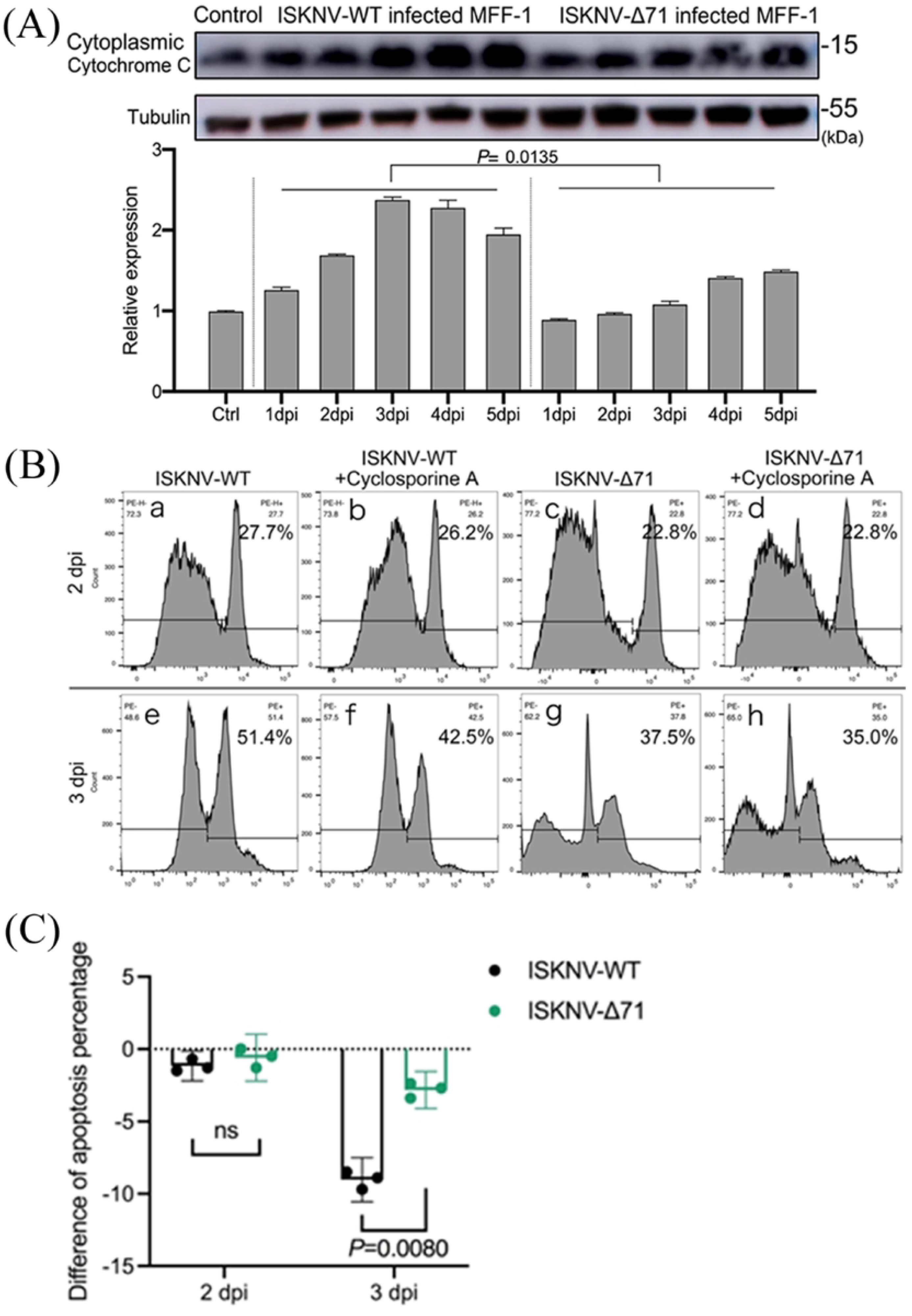
Mitochondrial cytochrome c (Cytc) release was higher in ISKNV-WT-infected cells than in ISKNV-△71-infected MFF-1 cells. (A) WB analysis of protein level of Cytc of MFF-1 cells after infected with ISKNV-WT or ISKNV-Δ71. Tubulin was used as the internal control. The protein expression was analyzed using ImageJ. As a result, in ISKNV-WT infection, the content of Cytc released from mitochondria to the cytoplasm increased with the infection time, reaching 2.5 times that of the control group at 3 dpi. In ISKNV-Δ71 infection, the release of Cytc also showed an upward trend, but the increase amplitude was lower than that in ISKNV-WT infection. The peak value appeared at 5 dpi, which was 1.5 times that of the control group. (B) Flow cytometry determined that CsA decreased the percentage of MFF-1 apoptosis induced by ISKNV-WT or ISKNV-Δ71. ISKNV-WT and ISKNV-Δ71 infected MFF-1 with MOI=0.1. At 1 hpi, CsA was added at a final concentration of 1 μM, and the apoptosis of MFF-1 was measured at 2 dpi and 3 dpi. (C) Statistical analysis of apoptosis percentage changes. At 3 dpi, CsA reduced the apoptosis induced by ISKNV-WT by 9% and that induced by ISKNV-Δ71 by 2.8%.

We used cyclosporin A (CsA), a mitochondrial permeability transition pore (MPTP) inhibitor (49–51), to block MPTP opening in MFF-1 cells and investigate its effect on apoptosis induced by ISKNV-WT and ISKNV-Δ71. CsA inhibits MPTP formation by blocking the interaction between CypD and ANT, thereby preventing conformational changes in ANT—both CypD and ANT have been identified as components of MPTP (52–55). Reported effective concentrations of CsA for MPTP inhibition in mammals range from 0.2 μM to 2 μM (49–51, 55, 56). We used the Cell Counting Kit-8 (CCK-8) to assess the toxicity of CsA to MFF-1 cells at these concentrations. The results showed that cell viability decreased in a CsA concentration-dependent manner at final concentrations of 0.1, 1, 5, and 10 μM. However, CsA had little effect on cellular respiratory activity at concentrations of 5 μM or lower (**Fig. 4C**).

A final concentration of 1 μM CsA was used to test its ability to inhibit apoptosis induced by ISKNV-WT. MFF-1 cells were infected with either virus strain at an MOI of 0.1; CsA was added 1 hpi, and cell apoptosis was detected at 2 and 3 dpi. At 2 dpi, the apoptosis rate induced by ISKNV-WT was 27.7%, which decreased to 26.2% in the CsA-treated group. For ISKNV-Δ71-infected cells, the apoptosis rate was 22.8% in both the CsA-untreated and treated groups. At 3 dpi, CsA significantly reduced the apoptosis rate induced by ISKNV-WT (51.4% in the untreated group vs. 42.5% in the treated group). For ISKNV-Δ71-infected cells, the apoptosis rate was 37.5% in the untreated group and 35% in the treated group (**Fig. 4B**). These results indicate that VP71 mediates apoptotic effects through an MPTP-dependent mechanism.

### VDAC2 is involved in the composition of ISKNV virions and interacts with p71

Based on viral proteomics data, five protein peptides perfectly matched VDAC2 of Japanese flounder (*Paralichthys olivaceus*) (**Fig. 5A**) in purified RSIV-type SKIV-ZJ07 (20), suggesting that VDAC2 may be involved in the assembly of mature *M. pagrus1* virions. In this study, we generated an antibody against *mf*VDAC2 (S2 Fig). Purified ISKNV was subjected to fractionation with TX-100 treatment, and WB analysis showed that *mf*VDAC2 was mainly detected in the supernatant fraction of TX-100-treated virus, while p71 was predominantly found in the pellet fraction (**Fig. 5B**). As controls, the envelope myristoylated membrane protein (MMP) p7 was mainly present in the supernatant, the DNA-binding protein (DBP) was exclusively in the pellet, and the MCP (p6) was detected in both fractions (**Fig. 5B**)—consistent with our previous findings on these proteins (19). Considering the mitochondrial localization of VDAC2 and its envelope-associated localization in the virus, we selected 16 viral structural proteins (p6, p7, p36, p37, p39, p41, p55, p56, p71, p77, p88, p93, p95, p101, p117, and p118) and 7 non-structural proteins (p2, p5, p25, p39, p84, p92, p102, p116) to investigate their potential interactions with *mf*VDAC2. Using *mf*VDAC2 as the bait protein (fused to the DNA-binding domain, BD) and the 23 viral proteins as prey proteins (fused to the DNA activation domain, AD), a yeast two-hybrid assay was performed to screen for potential interactions. As a result, only the co-transformation system of *mf*VDAC2-BD plus p71-AD yielded vigorously growing blue colonies on the final SD/-Leu/-Trp/-His/-Ade/X-α-gal quadruple dropout plate, indicating a potential interaction between *mf*VDAC2 and p71 (**Fig. 5C and D**). This interaction was further validated by co-immunoprecipitation (co-IP) assay (**Fig. 5E**).

**Fig. 5.**
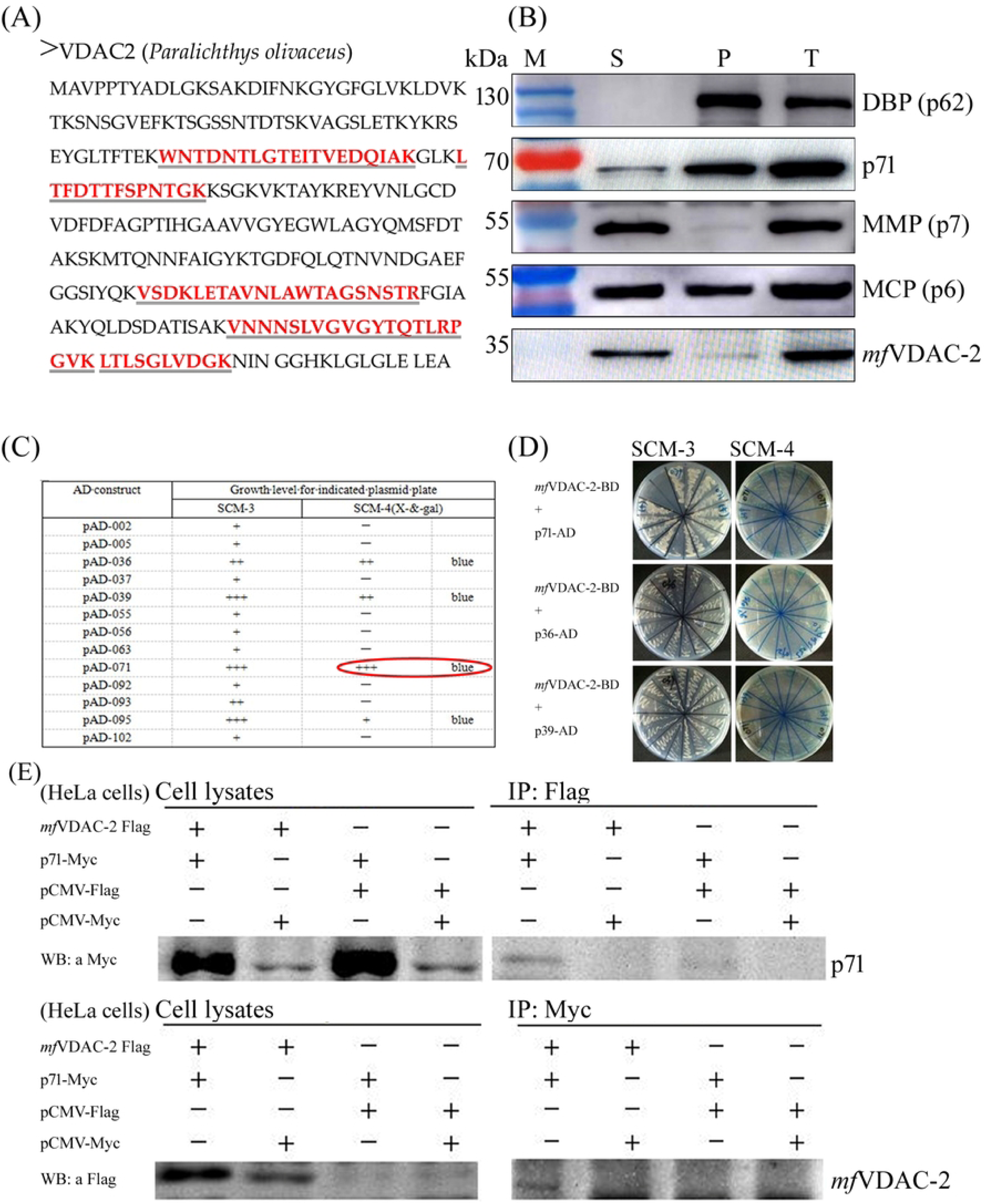
VDAC-2, as a host-associated protein, is incorporated into purified ISKNV/RISV virions and interact with p71. (A), Mass spectrometry identification of highly purified SKIV-ZJ07 virions showed that 5 amino acid sequences exactly matched the VDAC-2 protein of Japanese flounder (*P. olivaceus*); the 5 sequences with complete matches to Japanese flounder VDAC-2 are indicated by red bold double underlines. (B). WB analysis was performed to localize *mf*VDAC-2 in the structure of purified ISKNV virions, suggesting that *mf*VDAC-2 is localized to the envelope structure of purified ISKNV. S: Supernatant of TX-100-treated virions; P: Pellet of TX-100-treated virions; T: Total untreated viral proteins. DBP, DNA-binding protein (ISKNV-ORF062, p62), MMP, Myristoylated Membrane Protein (ISKNV-ORF007, p7); MCP, Major capsid protein (ISKNV-ORF006, p-6). (C, D), Partial result of Y2H in SCM3 and SCM4 agar indicates a potential robust interaction between *mf*VDAC2 and p71. (E), HeLa cells were co-transfected and the following combinations were collected: (1) *mf*VDAC2-Flag + p71-Myc; (2) *mf*VDAC2-Flag + Myc-tagged empty vector; (3) p71-Myc + Flag-tagged empty vector; (4) Flag-tagged empty vector + Myc-tagged empty vector. After lysis with IP lysis buffer, forward and reverse Co-IP assays were performed using anti-Flag and anti-Myc antibodies, respectively. WB results showed that *mf*VDAC2-Flag and p71-Myc could co-precipitate with each other, indicating that the interaction between *mf*VDAC2 and p71.

### *mf*VDAC2 is hijacked into the nucleus via interacting with nucleus-localized p71

Various recombinant plasmids encoding fused *mf*VDAC2 and p71 were constructed and transfected into MFF-1 (**Fig. 6A-C**), BHK21 (**Fig. 6D**), or HeLa (**Fig. 6E and F**) cells either individually or co-transfected. Both direct observation of fluorescent reporter genes or indirect immunofluorescence assay (IFA) results showed that *mf*VDAC2 and p71 were specifically localized to mitochondria and the nucleus of transfected cells, respectively (**Fig. 6A, B, E and F**). However, in cells co-transfected with both plasmids, obvious nuclear translocation of *mf*VDAC2 was observed (**Fig. 6C, D and F**). Notably, the intensity of *mf*VDAC2 nuclear translocation was positively correlated with the expression level of p71 in co-transfected cells (**Fig. 6D and F**). Co-IP) assays reconfirmed the interaction between *mf*VDAC2 and p71 (**Fig. 6G**), leading us to conclude that the nuclear translocation of *mf*VDAC2 is mediated by its interaction with nucleus-localized p71. MFF-1 cells were infected with ISKNV-WT or ISKNV-Δ71, and the subcellular localization of endogenous *mf*VDAC2 was traced by IFA using the anti-*mf*VDAC2 antibody. The results showed that endogenous VDAC2 underwent more widespread nuclear translocation in ISKNV-WT-infected MFF-1 cells compared to ISKNV-Δ71-infected cells (**Fig. 7A**). WB analysis also revealed a significant reduction in mitochondrial VDAC2 levels in ISKNV-WT-infected cells relative to ISKNV-Δ71-infected cells (**Fig. 7B**).

**Fig. 6.**
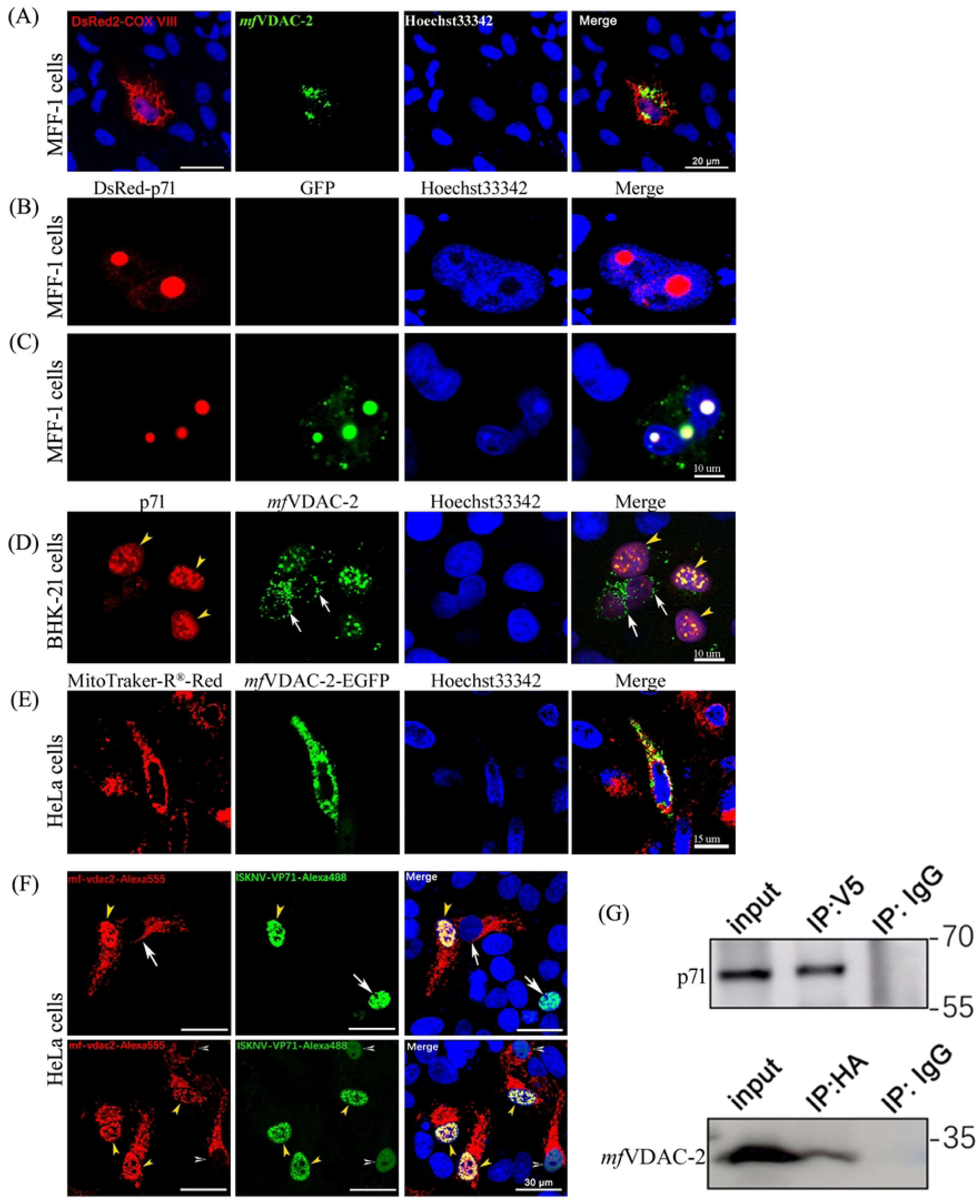
ISKNV-VP71 hijacks *mf*VDAC-2 from OMM into the nucleus via interacting with it. (A), EGFP-*mf*VDAC2 was transfected into MFF-1 cell, and mitochondrial probe DsRed2-cox-III and Hoechst 33342 were used for staining the cells. Confocal laser scanning imaging showed that *mf*VDAC2 is localized to the mitochondria of MFF-1 cells. Red fluorescence indicates the DsRed2 fluorescent protein fused with the mitochondria-targeting motif of cytochrome c oxidase subunit VIII (COX-VIII), which is localized to the inner mitochondrial membrane. Green fluorescence represents the EGFP-*mf*VDAC2, which is localized to OMM. (B), localization of p71 (red) by transferring recombinant DsRed-p71 in MFF-1 cells. The results showed that p71 is clear nucleus localization. (C), colocalization of *mf*VDAC2 and p71 by co-transferring DsRed-p71 and *mf*VDAC2-EGFP in MFF-1 indicates clear nucleus localization. (D) Colocalization of p71 and *mf*VDAC-2 in BHK-21 cells. *mf*VDAC2-Flag and p71-Myc were co-transfected into BHK21 cells. At 36 hpi, indirect immunofluorescence assay (IFA) was performed to detect the expression and localization of *mf*VDAC-2-Flag and p71-Myc. Mouse anti-Flag monoclonal antibody and rabbit anti-Myc polyclonal antibody were used as primary antibodies, while Alexa Fluor 488-conjugated goat anti-rabbit secondary antibody and Alexa Fluor 555-conjugated goat anti-mouse were used as secondary antibodies, respectively. Meanwhile, Hoechst 33342 was used to stain cell nuclei. The results showed that in cells co-transfected with *mf*VDAC-2-Flag and p71-Myc, *mf*VDAC-2 exhibited significant nuclear translocation (yellow arrowheads). In contrast, in cells without p71-Myc transfection or with low p71 expression, *mf*VDAC-2 (white arrows) was mainly localized in the cytoplasm (mitochondria). p71-Myc was consistently localized in the nucleus, suggesting that *mf*VDAC-2 is regulated to enter the nucleus through its interaction with p71. (E), localization of *mf*VDAC-2 (green) by transferring *mf*VDAC-2-EGFP into HeLa cells. Mitochondrial localization was traced by staining with Mito-Tracker^®^ Red (red). The results showed that *mf*VDAC-2 have clear mitochondrial localization in HeLa cell. (F) pcDNA3.1-V5-*mf*VDAC-2 vectors were co-transfected into HeLa cells. At 36 hours post-transfection, immunofluorescence analysis was performed to determine the subcellular localization of p71 and *mf*VDAC-2. Three expression patterns were observed: p71, when expressed alone, was localized in the nucleus (white arrows); *mf*VDAC-2, when expressed alone, was localized in the cytoplasm/mitochondrial (white arrows); when co-expressed, part of *mf*VDAC-2 co-localized with p71 in the nucleus and distributed in punctate aggregates (yellow arrowheads). This nuclear co-localization phenomenon was prevalent in co-transfected cells. When the expression level of p71 was low, the nuclear localization of *mf*VDAC-2 decreased accordingly (hollow arrowheads), suggesting that the nuclear localization of *mf*VDAC-2 is associated with the expression of p71. (G), Co-IP once again confirmed the interaction between VP71 and *mf*VDAC-2 (The tagged vectors pCMV-HA-ORF71 and pcDNA3.1-V5-*mf*VDAC-2 were co-transfected into HeLa cells).

**Fig. 7.**
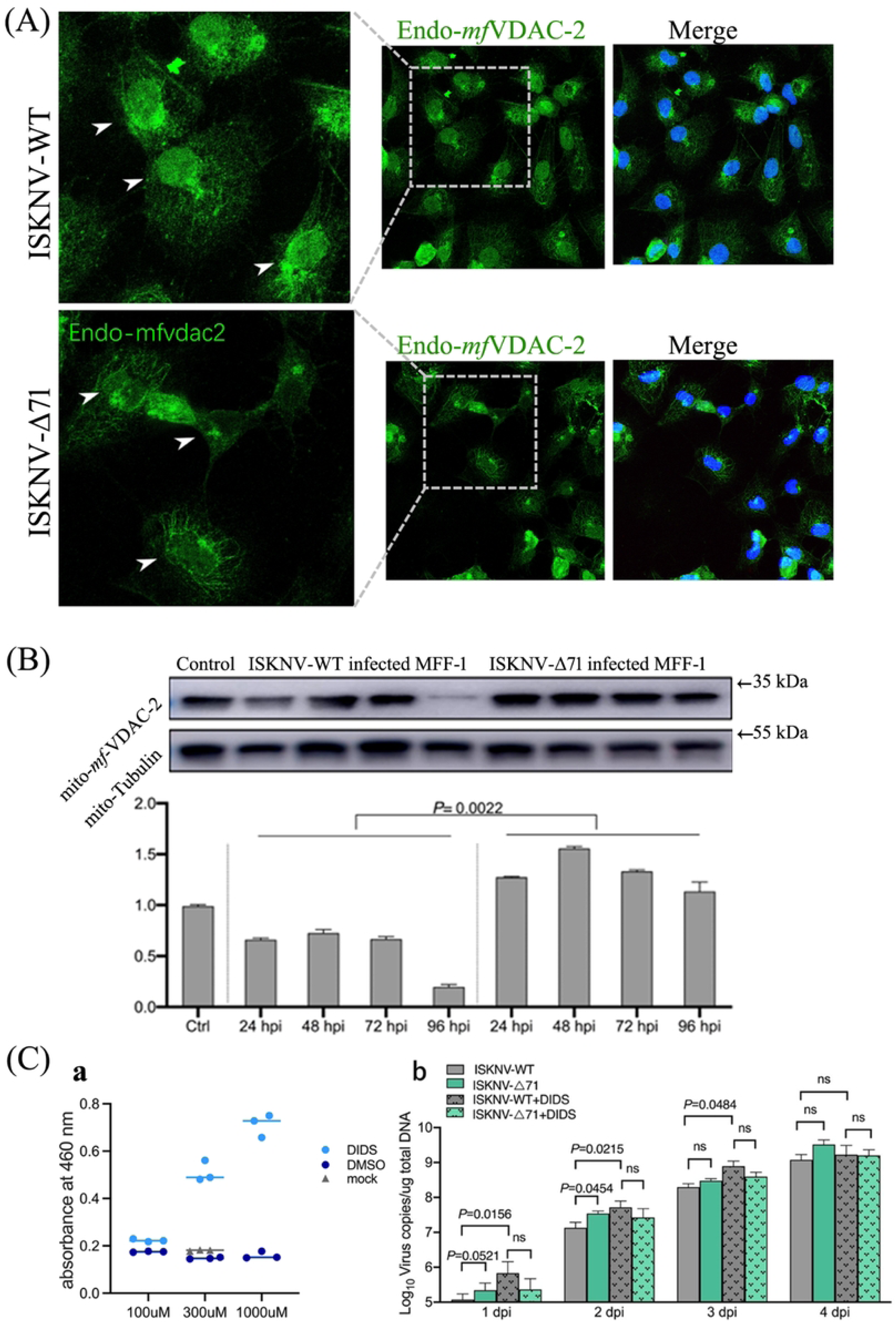
ISKNV-WT is more capable of inducing the nuclear translocation of endogenous *mf*VDAC2. (A), IFA analysis of endogenous *mf*VDAC2 localization. After MFF-1 cells were infected with ISKNV-WT or ISKNV-Δ71, the localization of endogenous VDAC2 protein was detected. Scale bar = 20 μm. (Panel a, b, c), Localization of endogenous *mf*VDAC2 in ISKNV-WT-infected MFF-1 cells; (Panel d, e, f), Localization of endogenous *mf*VDAC2 in ISKNV-Δ71-infected MFF-1 cells; (B), WB analysis of mitochondrial *mf*VDAC2 protein level. After MFF-1 cells were infected with ISKNV-WT or ISKNV-Δ71, the protein level of *mf*VDAC2 in mitochondria was detected. β-Tubulin was used as the internal control, and protein expression was analyzed by ImageJ software. In ISKNV-WT-infected cells, the level of mitochondrial *mf*VDAC2 continuously decreased, dropping to 20% of the control group at 96 hpi. In ISKNV-Δ71-infected cells, the relative expression level of mitochondrial *mf*VDAC2 was higher than that of the control group at all time points, which was consistent with the change trend of *mf*VDAC2 expression in whole cells. These results indicate that ISKNV-WT infection can downregulate the protein level of *mf*VDAC2 in mitochondria of MFF-1 cells, and this downregulation is mediated by VP71. (**C),** Effect of VDAC inhibitor on ISKNV/ISKNV-Δ71 infection. (a) Effect of DIDS on cellular respiratory activity detected by CCK-8 assay: The x-axis represents the final concentration of DIDS, and the y-axis represents the absorbance at 460 nm. An increase in absorbance indicates enhanced respiratory activity of MFF-1 cells. (b) Effect of DIDS on the replication of ISKNV-WT and ISKNV-Δ71: Statistical analysis was performed using the t-test, and ns indicates no statistically significant difference.

Given the association between VDAC2 and apoptosis (57), a comparative analysis of ISKNV-WT and ISKNV-Δ71 infection groups was performed using the apoptosis inhibitor DIDS. Cells were seeded at the same density in culture dishes and treated with DIDS at final concentrations of 100 μM, 300 μM, and 1000 μM, with the same volume of DMSO as the negative control and untreated cells as the blank control. After 24 hours of incubation, cell metabolic activity was measured using the CCK-8 assay. The results showed that cell metabolic activity gradually increased with increasing DIDS concentration, and DIDS at a final concentration of 100 μM had little effect on cell growth and metabolism (**Fig. 7C-a**). qPCR analysis showed no significant difference in viral copy numbers between ISKNV-Δ71 and ISKNV-WT in the DIDS-treated groups, while the copy number of ISKNV-Δ71 was higher than that of ISKNV-WT in the untreated group (**Fig. 7C-b**), suggesting that the slower *in vitro* replication rate (21) of ISKNV-WT compared to ISKNV-Δ71 may be caused by p71-mediated apoptosis.

### Zebrafish VDAC2 also interacts with p71 and is induced to translocate into the nucleus

We analyzed the evolutionary homology between *mfvdac2* and three zebrafish *vdac* isoforms (*zfvdac1, zfvdac2, zfvdac3*). The results showed high homology between *mf*VDAC2 and the three *zf*VDACs (**Fig. 8A-a**). Specifically, *mf*VDAC2 shared 93.99% homology with *zf*VDAC2, 75.62% and 80.5% amino acid sequence homology with *zf*VDAC1 and *zf*VDAC3, respectively. Considering conservative amino acid substitutions, the sequence similarity increased to 92.93% and 89.72% (**Fig. 8A-b**). Subcellular localization and co-localization assays demonstrated that all three *zf*VDACs were specifically localized to the cytoplasm/mitochondria (**Fig. 8B and D**). However, upon co-transfection with p71, only *zf*VDAC2 underwent significant nuclear translocation. Notably, the intensity of *zf*VDAC2 nuclear translocation was positively correlated with the expression level of p71 in co-transfected cells (**Fig. 8B and D**). These results are highly consistent with the observed subcellular interaction between *mf*VDAC2 and p71 (**Fig. 6**). Co-IP results also confirmed the interaction between p71 and *zf*VDAC2 (**Fig. 8C**).

**Fig. 8.**
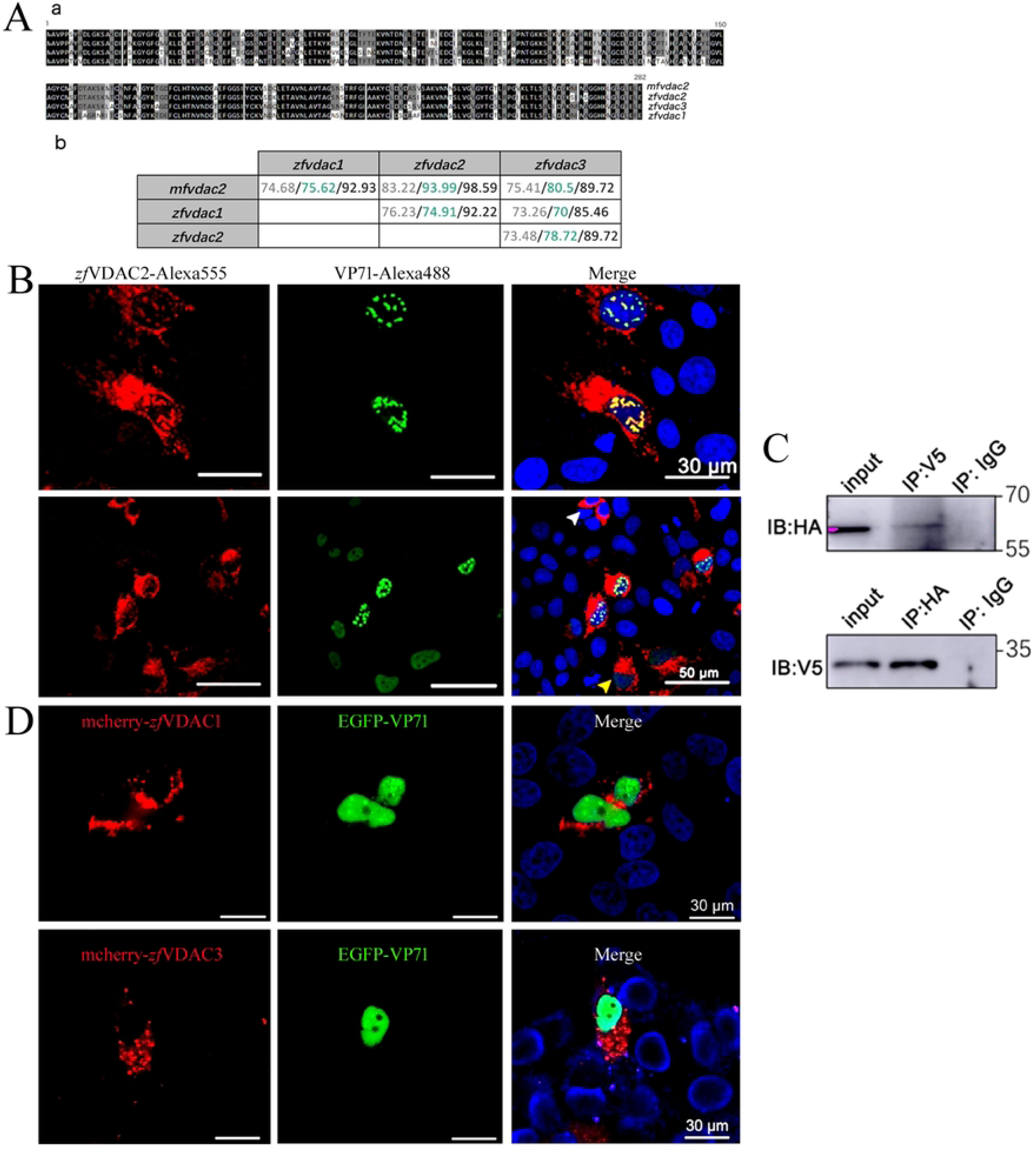
Zebrafish VDAC2 (*zf*VDAC2) is highly homologous to *mf*VDAC2), and can be also hijacked into the nucleus by p71. (A), Multiple sequence alignment of the homology between *mf*VDAC2 and zebrafish VDACs; (B), p71 can induce the nuclear translocation of *zf*VDAC2. HeLa cells were co-transfected with the vectors pCMV-HA-ORF71 and pcDNA3.1-V5-zfVDAC2, and IFA was performed at 36 hours post-transfection as the above mentioned (Fig. 6F). (C), Co-IP assay confirmed the interaction between *zf*VDAC2 and p71; (D), *zf*VDAC1 and *zf*VDAC3 do not co-localize with p71. pmCherry-N1-*zf*VDAC1 and pmCherry-N1-*zf*VDAC3 were co-transfected with pEGFP-N1-ISKNV ORF71 into HeLa cells respectively, and the subcellular localization of the proteins was observed under a confocal microscope.

### VDAC2 knockout zebrafish does not significantly affect the pathogenicity of ISKNV-WT and ISKNV-Δ71

Using CRISPR/Cas9 gene edition technology, we successfully generated a *vdac2* knockout zebrafish line *(zfvdac2^-/-^*). The zebrafish *vdac2* gene (Ensembl: ENSDARG00000013623) is located on chromosome 13, with a full length of 15,178 bp. It contains 9 exons, with the start codon ATG in exon 2 and the stop codon TAA in exon 9 (**Fig. 9A-a**). A sgRNA targeting the 3rd coding exon was designed based on Cas9 cleavage rules and off-target prediction. The gRNA and Cas9 mRNA were injected into zebrafish embryos, and F0 generation *zfvdac2* knockout individuals were screened at 3 months of age by extracting DNA from tail fins. Gene editing efficiency was analyzed by PCR amplifying the 3rd exon sequence followed by Sanger sequencing of the PCR products. Specifically, sequencing electropherograms were examined; when double peaks appeared, the number of deleted or inserted bases was identified by analyzing the phase difference of the overlapping peaks (**Fig. 9B**). Finally, one heterozygous male zebrafish with a 7 bp deletion (*vdac2^+/-7^*) (**Fig. 9Ba**) and one heterozygous female zebrafish with a 10 bp deletion (*vdac2^+/-10^*) (**Fig. 9Bb**) were identified, designated as the F0 generation.

**Fig. 9.**
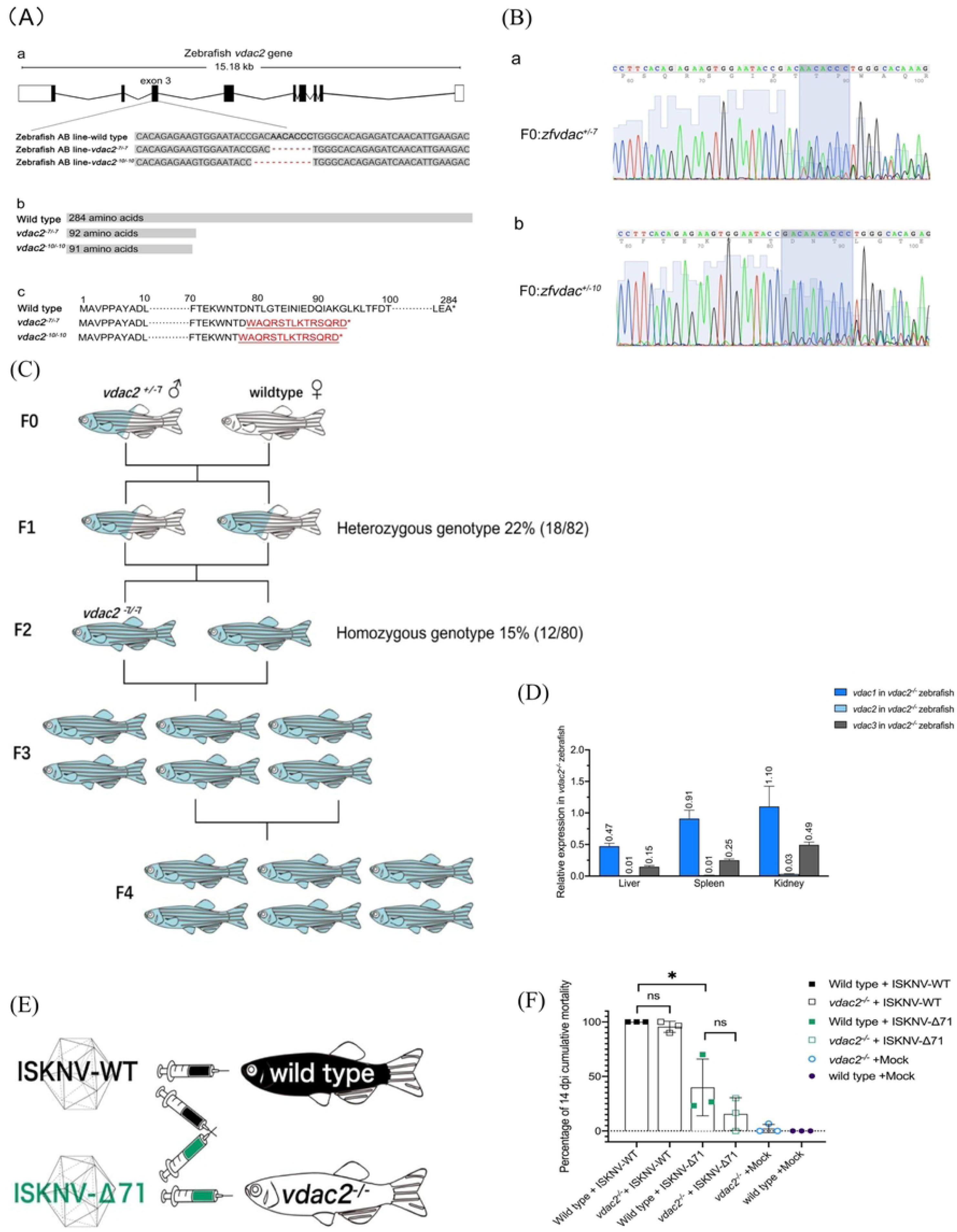
Establishment and characterization of the *vdac2* knockout zebrafish line. (A), Gene structure and knockout site of zf*vdac2*, gene editing leads to the premature termination codon (PTC), which leads a truncated protein. (a) Schematic diagram of the structure and knockout site of the zebrafish *vdac2* gene. (b) Gene editing results in the premature appearance of a stop codon, leading to the translation of a truncated protein. (c) Frameshift mutation results in the translation of an aberrant amino acid sequence. (B), Electropherograms of PCR - amplified exon 3 of the *zfvdac2* gene in F0 generation zebrafish. (a) The double sequencing peaks induced by a 7 - bp base deletion are marked with a blue box. (b) The double sequencing peaks induced by a 10 - bp base deletion are marked with a blue box. (C), Screening and breeding flow chart of *zfvdac2* gene knockout zebrafish. (D), qPCR analysis of zf*vdac1*, zf*vdac2*, zf*vdac3* transcription in *vdac2^-/-^* zebrafish liver, spleen and kidney relative to that of wild type zebrafish; (E), Schematic diagram of the four challenge combinations in the zebrafish challenge experiment; (F), Statistical analysis of cumulative mortality in zebrafish challenge experiment at 14 dpi. Using *t*-test method to statistical analysis. * P < 0.05. ns, not significant.

The screening process is illustrated using the example of obtaining homozygous zebrafish with a 7 bp deletion in the 3rd exon of the *zfvdac2* gene (*zfvdac2^-7/-7^*) (**Fig. 9Ab, Ac**). Crossing the F0 heterozygous male zebrafish (*vdac2^+/-7^*) with wild-type female zebrafish yielded F1 generation zebrafish with either wild-type or *vdac2^+/-7^* genotypes. At 3 months of age, DNA was extracted from the tail fins of F1 zebrafish, and PCR analysis of the 3rd exon of the *zfvdac2* gene was performed. Heterozygous *vdac2^+/-7^* individuals accounted for 22% (18/82 individuals) (**Fig. 9C**).

Reciprocal challenge experiments were performed on wild-type zebrafish and *zfvdac2^-/-^* zebrafish with ISKNV-WT and ISKNV-Δ71 (**Fig. 9E**). The results showed that ISKNV-WT exhibited high lethality to both wild-type and *zfvdac2^-/-^* zebrafish, while ISKNV-Δ71 showed significantly attenuated lethality in both zebrafish strains (**Fig. 9F and S3 Fig**). Histopathological analysis revealed that, unlike in mandarin fish spleen infected with ISKNV, characteristic swollen cells of ISKNV infection were rarely observed in the tissues of all four groups of zebrafish—consistent with previous reports (58). Additionally, necrotic lesions were reduced and tissue compactness was increased. Abundant red blood cells were observed in the spleen tissues of both zebrafish strains infected with ISKNV-Δ71, reflecting good tissue perfusion (**Fig. 10A-a, -b, -e, -f)**. In contrast, red blood cells were reduced in ISKNV-WT-infected spleen tissues, which may indicate more severe tissue damage (**Fig. 10A-c, -d, -g, -h**). However, no significant differences were observed between the two zebrafish strains infected with the same virus.

**Fig. 10.**
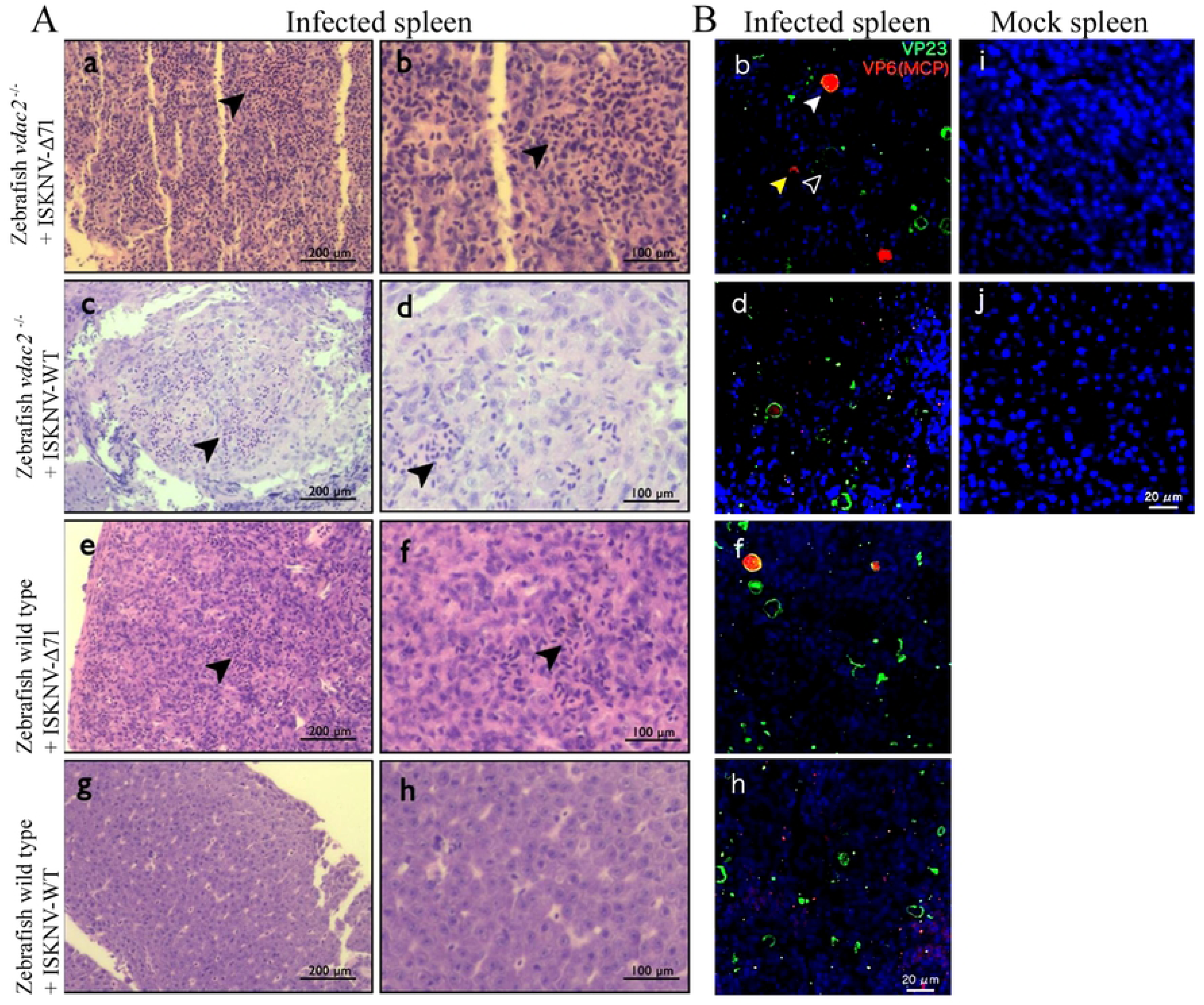
Histopathological and immunofluorescence analyses of the cross-infections of ISKNV-WT or ISKNV-Δ71 in *zfvdac2* or *zfvdac2^⁻/⁻^* zebrafish. A, Tissue sections of zebrafish infected with ISKNV, H&E staining analysis. Sampling at 4 dpi. The image was taken under an upright brightfield microscope with a magnification of 40×. Panels a, c, e, and g show the pathological findings of the two ISKNV strains infecting the two types of zebrafish, while panels b, d, f, and h are the corresponding 2× magnified images. Black arrows indicate red blood cells. (B), Immunofluorescence analysis of zebrafish tissue sections. ISKNV-VP23 is labeled as green fluorescence, and ISKNV-MCP is labeled as red fluorescence. Solid white arrows indicate infected cells recognized by both VP23 and MCP, along with a large number of virions; open arrows indicate infected cells recognized only by VP23; yellow arrows indicate cells labeled only by MCP, which contain a large number of mature virus particles. Panels b, d, f and h indicate the pathological findings of the two ISKNV strains infecting the two types of zebrafish. Panels i and j indicate the mock infection spleen.

Immunofluorescence analysis showed that typical swollen cells—characteristic markers of ISKNV infection—were present in the spleen tissues of all four challenge groups (**Fig. 10B**). As indicated by solid white arrows, swollen cells filled with MCP were surrounded by basement membranes containing VP23; open arrows indicate ruptured basement membranes with released virions, resulting in undetectable MCP in the cells. Yellow arrows indicate red fluorescence representing virions released into the tissues. Immunofluorescence results demonstrated extensive viral release into the tissues in the spleens of both zebrafish strains infected with ISKNV-WT at 4 dpi (**Fig. 10B-d, -h**), which was significantly higher than that in the ISKNV-Δ71 groups (**Fig. 10B-b, -f**). No fluorescent signals were detected in the mock group, excluding false positives caused by non-specific antibody binding (**Fig. 10B-i, -j**). No differences were observed between the two zebrafish strains infected with ISKNV-Δ71, nor between those infected with ISKNV-WT, in the immunofluorescence analysis.

### Knockout of *zfvdac2* does not induce a genetic compensation response

The above results indicate that *zfvdac2* knockout does not affect the replication and virulence of ISKNV-Δ71. We hypothesized whether *zfvdac2* knockout activates a genetic compensation response, leading to increased expression levels of its homologous genes *zfvdac1* and *zfvdac3* to compensate for and replace the function of *zfvdac2*. Studies have identified a genetic compensation mechanism in zebrafish, wherein knockout of a gene can upregulate the expression of its homologous genes for genetic compensation(59, 60). The molecular mechanism of this genetic compensation response has been elucidated: certain base mutations can result in the transcription of nonsense mRNAs (containing premature termination codons, PTCs). These nonsense mRNAs are degraded via the nonsense-mediated mRNA decay (NMD) pathway to prevent the translation of aberrant proteins. Upf3a, a homologous protein of Upf3b that mediates NMD, can interact with nonsense mRNAs and the COMPASS (complex of proteins associated with Set1) complex. It guides the complex to the coding regions of homologous genes through nonsense mRNAs, thereby promoting the expression of homologous genes by methylating histones near the coding regions (61, 62). Our analysis revealed that the *zfvdac2* gene in *zfvdac2^-/-^* zebrafish contains multiple premature termination codons (PTCs) following the deletion of 7 codons, which is consistent with the recognition mechanism of the genetic compensation response. Therefore, we measured the transcriptional changes of *zfvdac1*, *zfvdac2*, and *zfvdac3* at the *in vivo* level to explore whether *zfvdac2* knockout affects their transcription. Total RNA was extracted from the liver, spleen, and kidney of 10 healthy *zfvdac2^-/-^* zebrafish and 10 wild-type zebrafish, followed by reverse transcription into cDNA. Relative quantitative PCR (qPCR) analysis was performed using the β-actin gene as an internal reference. The results showed (**Fig. 9D**) that the transcriptional level of *zfvdac1* in the liver of *zfvdac2^-/-^* zebrafish was 47% of that in wild-type zebrafish, with no significant changes in the spleen and kidney. The transcriptional level of *zfvdac2* mRNA was drastically reduced in the liver, spleen, and kidney tissues of *zfvdac2^-/-^* zebrafish, accounting for only 1%-3% of that in wild-type zebrafish—indicating that the *zfvdac2* gene may be degraded via the NMD pathway. Unexpectedly, *zfvdac3* mRNA was downregulated: its transcriptional level in the liver of *zfvdac2^-/-^* zebrafish was 15% of that in wild-type zebrafish, while in the spleen and kidney, the values were 25% and 49%, respectively. These results demonstrate that *zfvdac2* knockout does not induce a genetic compensation response characterized by increased transcription of *zfvdac1*/*zfvdac3*, and the downregulation of *zfvdac3* transcription requires further confirmation.

## Discussion

The fates of cells upon ISKNV/RSIV infection, such as apoptosis, necrosis, or autophagy, have been briefly described; however, most studies merely focus on phenotypic observations and lack in-depth mechanistic analysis (17, 37, 39, 40, 63). We here found that infection of MFF-1 cells with ISKNV-WT induces higher levels of apoptosis compared to infection with ISKNV-Δ71. To first identify the pathway through which VP71 mediates apoptosis, we verified that ΔΨm was decreased in MFF-1 cells infected with either ISKNV-WT or ISKNV-Δ71 relative to the control group. However, the ΔΨm was significantly lower in ISKNV-WT-infected cells than in ISKNV-Δ71-infected cells, indicating more severe mitochondrial dysfunction induced by ISKNV-WT infection. Subsequently, we confirmed that this reduction in membrane potential was caused by increased mitochondrial membrane permeability. TEM observations also revealed that mitochondrial integrity was lower in ISKNV-WT-infected cells compared to ISKNV-Δ71-infected cells, with more prominent mitochondrial swelling and rupture observed in ISKNV-WT-infected cells. This suggests that mitochondrial integrity disruption may be triggered by the opening of MPTP. Further experiments using CsA to inhibit MPTP showed that ISKNV-WT-induced apoptosis was partially inhibited, demonstrating that VP71 indeed promotes MPTP-dependent apoptosis. However, CsA did not reduce the apoptosis level induced by ISKNV-WT to that observed in ISKNV-Δ71-infected cells, indicating that VP71 exerts its pro-apoptotic function not only through the MPTP pathway but also potentially via other mechanisms. Following apoptosis inhibition with DIDS, the replication level of ISKNV-WT was comparable to that of ISKNV-Δ71. This suggests that the slower replication of ISKNV-WT *in vitro* relative to ISKNV-Δ71 is due to its induction of stronger apoptosis.

VDAC2 was previously identified to be involved in the packaging of RSIV-type *M. pagrus1* virions (20), but its specific function has not been further elaborated. In this study, we generated a polyclonal antibody against *mf*VDAC2, and finally verified the localization of *mf*VDAC2 in the viral envelope via WB analysis. Using *mf*VDAC2 as the bait protein, we selected 23 ISKNV proteins as prey proteins and performed a Y2H assay to screen for potential viral interactors with *mf*VDAC2. Notably, considering the mitochondrial localization of VDAC2, our selected viral proteins included all 10 virus-encoded proteins predicted to be mitochondrially localized. Meanwhile, given the envelope localization of *mf*VDAC2 in ISKNV virions, the prey proteins also included 3 known viral envelope proteins (p7, p56, and p118) (19). As the most abundant structural protein (19, 20), the major capsid protein (MCP) often plays a crucial role in mediating iridovirus infection of hosts (38, 64) and was therefore naturally included as a potential interactor. Unexpectedly, the Y2H screen showed that *mf*VDAC2 only specifically interacted strongly with VP71, which was further validated by co-IP assay.

Of particular note, subcellular localization assays revealed that *mf*VDAC2 and VP71 were specifically localized to mitochondria and the nucleus, respectively, when expressed individually. However, in cells co-transfected with both *mf*VDAC2 and VP71, obvious nuclear translocation of *mf*VDAC2 was observed, and the extent of *mf*VDAC2 nuclear translocation was positively correlated with the expression level of VP71. This result reconfirms the strong interaction between *mf*VDAC2 and VP71, in which VP71 appears to play a dominant role, and VDAC2 translocates into the nucleus passively. Studies have indicated the involvement of VDAC2 in viral infection and pathogenicity: for example, VDAC2 was identified as a functional receptor mediating the entry and infection of lymphocystis disease virus (LCDV), another well-known fish iridovirus, into host cells, though the viral protein interacting with VDAC2 has not been uncovered (44). In studies on avian infectious bursal disease virus (IBDV), VDAC2 was shown to interact with the viral protein VP5 to induce apoptosis in infected cells—a process considered an active host defense mechanism that inhibits viral replication via VDAC2-mediated apoptosis (57). However, no translocation of VDAC2 from the OMM to the nucleus has been reported in either LCDV or IBDV infections. This study demonstrates that ISKNV infection manipulates the nuclear translocation of VDAC2 via interacting with VP71, thereby enhancing apoptosis in infected cells to a certain extent—representing a novel discovery of VDAC2 function. Notably, knockout of VP71 reduced apoptosis in infected cells to some degree but did not completely eliminate it, indicating that VP71-induced apoptosis via hijacking VDAC2 into the nucleus is just one component of viral-induced cell death, and may not be the major contributor. Megalocytiviruses represented by ISKNV are large DNA viruses with large genomes, encoding numerous proteins and exhibiting complex pathogenic mechanisms. Several forms of cell death, including apoptosis, necrosis, autophagy, and even ferroptosis, have been implicated in their infection (37–40, 63). Further in-depth studies are needed to unravel the intricate virus-host interactions.

Previous studies have reported that deletion of VDAC2 renders the mitochondrial apoptotic pathway more sensitive to apoptotic stimuli (25). Consistent with this, mitochondrial VDAC2 levels were lower in ISKNV-WT-infected MFF-1 cells than in ISKNV-Δ71-infected cells, accompanied by a higher apoptosis rate. Thus, we hypothesize that the reduction in mitochondrial VDAC2 is one of the mechanisms underlying the high apoptosis rate induced by ISKNV-WT infection. Based on these observations, we propose a model (**Fig. 11**) wherein VP71 induces the nuclear translocation of *mf*VDAC2, leading to a decrease in mitochondrial VDAC2 levels in MFF-1 cells. This, in turn, enhances the sensitivity of cells to ISKNV-WT-induced apoptosis, and the increased apoptosis ultimately slows down ISKNV replication. During TEM observations, we noted more severe mitochondrial swelling in ISKNV-WT-infected cells. Mitochondrial swelling is one of the morphological hallmarks of mitochondrial permeability transition (MPT); however, it is important to note that mitochondrial swelling can also be induced by oxidative stress (65, 66) or calcium overload (67). Further studies are required to determine whether VP71 mediates mitochondrial swelling and mitochondrial membrane permeabilization through these additional pathways.

**Fig. 11.**
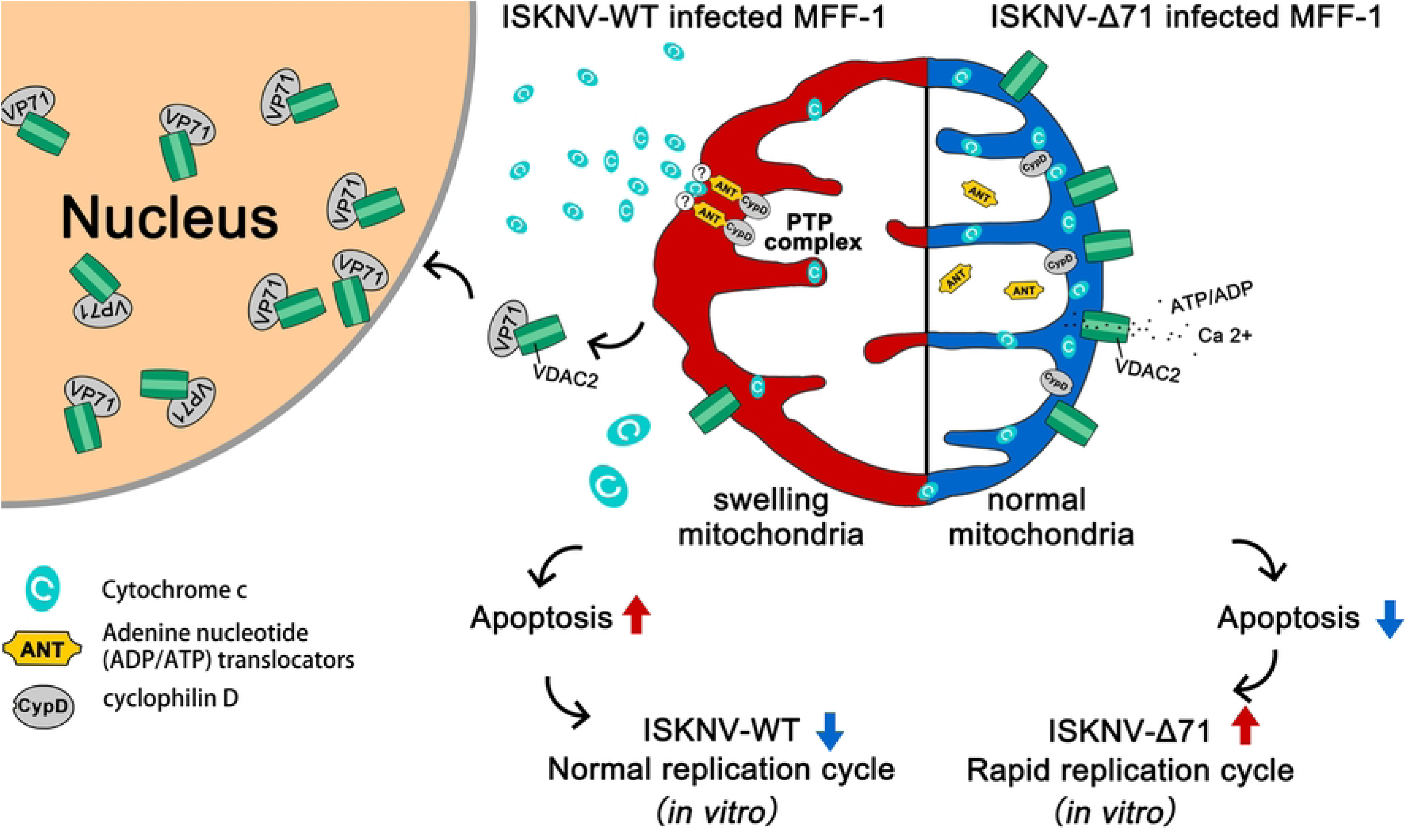
Schematic diagram illustrating the mechanism by which ISKNV-p71 hijacks VDAC2 from OMM to the nucleus, then enhance apoptosis by disrupting mitochondrial membrane permeability. In ISKNV-Δ71 infected MFF-1, *mf*VDAC2 localizes to mitochondria and inhibits the opening of MPTP, leading to a reduced apoptosis, then ISKNV-Δ71 replicates faster. In ISKNV-WT infection, *mf*VDAC2 interacts with VP71 to enter the nucleus, resulting in the decrease of mitochondrial *mf*VDAC2, the MPTP open leading an increased apoptosis, then slowed down the replication of ISKNV-WT.

VDAC2 is responsible for the transport of ATP/ADP and regulates mitochondrial respiratory rate to adapt to intracellular energy demands (68). We therefore investigated whether VDAC2 modulates viral replication rate by altering respiratory rate, thereby optimizing viral survival in the host. The CCK-8 assay was used to measure the respiratory activity of MFF-1 cells infected with ISKNV-WT or ISKNV-Δ71. No significant difference in respiratory activity was observed between the two viral strains at 6 and 12 hpi. However, at 18, 24, and 48 hpi, the respiratory activity of ISKNV-Δ71-infected cells was higher than that of ISKNV-WT-infected cells. These results require further validation using additional respiratory activity detection methods.

Apoptosis is widely recognized as a key host defense mechanism against viral infections. For example, apoptosis of virus-infected cells can recruit macrophages for phagocytosis and elimination of the infection (69), which appears contradictory to the pro-apoptotic role of ISKNV VP71. Previous studies have shown that ISKNV has evolved strategies to evade direct killing by immune cells. For instance, the viral-encoded protein VP23 can participate in the formation of a mock-basement membrane and recruit the adhesion of lymphatic endothelial cells, which may isolate immune cells and prevent direct killing (13). From this perspective, ISKNV may not need to inhibit apoptosis to evade direct immune cell-mediated clearance. In contrast, inducing apoptotic cell death in the late stage of infection may reduce the release of inflammatory mediators and attenuate the inflammatory response, which is beneficial for viral propagation. Many viruses encode pro-apoptotic proteins, indicating that inducing apoptosis at certain stages of the viral life cycle may be advantageous for viral survival. Although ISKNV-Δ71 induces lower apoptosis and exhibits faster replication *in vitro*, we observed attenuated virulence in infected mandarin fish. This seemingly contradictory result may be attributed to the fact that i*n vitro* viral infection experiments only partially simulate *in vivo* conditions. Viral replication in the natural host is subject to numerous constraints, such as the pressure to release progeny viruses from primary infection sites and achieve systemic dissemination, the pressure from the host immune system, and the pressure to efficiently transmit between hosts. We hypothesize that the reduced apoptosis caused by VP71 deleted ISKNV-Δ71 facilitates viral replication i*n vitro*, possibly due to enhanced mitochondrial integrity and higher respiratory rate accelerating the viral infection process. In contrast, VP71 deletion in ISKNV-Δ71 may upregulate cxcl12a or other cytokines during *in vivo* infection, which could stimulate and strengthen the host immune response, leading to attenuated virulence and reduced replication in mandarin fish (21). The ultimate goal of all viruses is to infect target cells, replicate large numbers of progeny virions, and transmit these progenies to initiate new rounds of infection. If triggering apoptosis were detrimental to viral replication, viruses would likely have evolved mechanisms to evade this process. Therefore, we speculate that the pro-apoptotic protein encoded by ISKNV-WT may be beneficial for viral survival in the natural host, representing a trade-off strategy for the virus to balance replication speed with other survival pressures.

VDAC2 is highly conserved (70). Alignment of the amino acid sequence of *mf*VDAC2 with that of zebrafish VDAC2 (*zf*VDAC2) revealed a 94% amino acid sequence similarity. When considering conservative substitutions, the homology reaches as high as 98%. We confirmed the interaction between VP71 and *zf*VDAC2 through co-IP and subcellular co-localization assays. Given the immature CRISPR/Cas9 gene editing technology for mandarin fish and zebrafish being a valid infection model for ISKNV (58), we therefore transferred the *in vivo* interaction model of VDAC2 and p71 to zebrafish. We attempted to construct a research model by knocking out the *zfvdac2* gene in zebrafish. Knockout of *vdac2* has been reported in mice: *vdac2*-knockout embryos fail to survive (30). However, another study showed that *vdac2*-knockout mouse embryos can be born alive, though most individuals die within three weeks after birth, with the maximum survival time of 40 days (71). In addition, knockout of *vdac1* in mice increases embryonic lethality and reduces live births, while *vdac3*-knockout male mice are infertile (25, 72). These findings indicate that mouse VDAC2 has non-redundant functions. Thus, *zfvdac2* knockout can not only be used to study the role of *zf*VDAC2 in ISKNV infection but also serve as a model for investigating the functions of *zf*VDAC2. Researchers have used *vdac2^-/-^* zebrafish to study the role of *zf*VDAC2 in Ca^2+^ uptake in cardiomyocytes and cardiac rhythm regulation (62, 73), which demonstrates that *vdac2*-knockout zebrafish can survive and reproduce. However, the phenotype of *zfvdac2*^-/-^ was not reported in that paper.

In this study, we observed that *vdac2* knockout induces a lethal effect. For instance, self-crossing of F1 *vdac2^+/-7^* individuals produced the F2 generation. Among F2 embryos, 55% of fertilized eggs developed to adulthood. At 3 months of age, DNA was extracted from the tail fins of F2 zebrafish for PCR analysis of the 3rd exon of the *zfvdac2* gene. Homozygous *zfvdac2* knockout individuals (*vdac2^-7/-7^*) accounted for 15% (12/80 individuals), including 5 males and 7 females. The genetic ratio of homozygous individuals was lower than the theoretically predicted 25%. Self-crossing of F2 *vdac2^-7/-7^* individuals yielded a large number of *vdac2^-7/-7^* individuals in the F3 generation. F3 zebrafish were used for viral challenge experiments after reaching adulthood. Self-crossing of F3 *vdac2^-7/-7^* individuals produced a large number of *vdac2^-7/-7^* individuals in the F4 generation, which were used for statistics on the natural mortality rate and viral challenge experiments of *vdac2^-7/-7^* zebrafish. *vdac2^-7/-7^* zebrafish were selected for subsequent experiments, and are referred to as *vdac2^-/-^* in the following text. During the self-crossing of F2 *vdac2^-/-^*individuals to breed the F3 generation, an abnormally high embryonic mortality rate was observed. Additionally, 58.3% (7/12) of F2 *vdac2^-/-^* parent fish died by 5 months postfertilization (mpf), indicating that defects caused by *vdac2* knockout lead to high mortality. To detail the mortality rate, 4 groups of F3 zebrafish (each containing 2 males and 1 female) were selected. Fertilized eggs were collected separately after natural spawning, with a total of 1,147 fertilized eggs subjected to mortality statistics over 300 days. A total of 211 fertilized eggs from wild-type zebrafish were included for the same statistics.

In the overall mortality rate graph (S4 Fig. A), two periods of rapid survival rate decline were observed: the first at 24 hours postfertilization (hpf), and the second from 5 to 7 dpf. Furthermore, a constant mortality rate was observed in the *vdac2^-/-^* zebrafish population after approximately 3 mpf; only 14.65% of *vdac2^-/-^* zebrafish survived by 300 dpf, compared to 98.1% survival in wild-type zebrafish (S5 Fig). These results demonstrate that *zf*VDAC2 is an essential functional gene in zebrafish.

Observation of dying larvae at 6 dpf under a stereomicroscope revealed delayed development, with significantly smaller yolk sacs and swim bladders compared to surviving *vdac2^-/-^* individuals. No visible phenotypic differences were observed between surviving *vdac2^-/-^* individuals and wild-type individuals (S4 Fig. B). The lethal mechanisms underlying the mortality peaks at 24 hpf and 4-7 dpf may differ. Embryos develop in a sterile environment within the chorion before 3-4 dpf; mortality at 24 hpf may be caused by excessive apoptosis leading to developmental disorders (25, 74). Larval intestines are colonized by microorganisms 12-24 hours after chorion rupture (75, 76), so mortality during this period may result from bacterial infection or defects in oocytes due to parental reproductive system abnormalities (77). The specific mechanisms require further investigation.

To more precisely describe the subsequent mortality rate, the survival rate from 7 dpf to 300 dpf was calculated using the number of surviving individuals at 7 dpf as the baseline (set to 100% survival rate) (S6 Fig). All *vdac2^-/-^* individuals surviving beyond 7 dpf were capable of swimming and feeding. Twenty *vdac2^-/-^* zebrafish larvae died between 7 dpf and 14 dpf, accounting for 5.6% of the total 358 individuals.

Notably, the mortality rate of *vdac2^-/-^* zebrafish decreased sharply after 14 dpf: only one individual died at 15 dpf and 16 dpf, respectively, with no deaths from 17 dpf to 89 dpf. The mortality rate began to gradually increase after 90 dpf, slowing slightly around 240 dpf. From 8 dpf (when fry is sufficiently developed to feed) to 300 dpf, 53.1% of *vdac2^-/-^* individuals died. In contrast, the survival rate of wild-type zebrafish remained at 100% until the first death occurred at 235 dpf, with a survival rate of 98.58% at 300 dpf (S6 Fig). These results indicate that *vdac2* knockout in zebrafish causes defects leading to massive deaths from the gastrula stage to early larval stage. No deaths of *vdac2* knockout individuals occurred between 17 dpf and 90 dpf. The survival rate of *vdac^2-/-^* zebrafish began to decline again after 90 dpf, a trend that continued until 300 dpf. This suggests that the mechanism causing death in adult zebrafish may differ from that responsible for the early mortality peak (at 24 hpf).

To investigate the cause of abnormal deaths in adult *vdac2^-/-^* zebrafish, 25 *vdac2^-/-^* individuals with behavioral or morphological abnormalities were observed. Their symptoms mainly included: 100% (25/25) showing isolation, cessation of feeding, and slow swimming; 8% (2/25) with scale loss; 68% (17/25) with redness and swelling in the entire abdomen or posterior abdomen; and 20% (5/25) with emaciation. Dissection of diseased fish revealed symptoms such as swollen digestive tracts, liver hemorrhage, ascites, and ovarian ulceration in females (S4 Fig. C). For *vdac2^-/-^* zebrafish at 6 mpf, sampling was conducted at approximately one-week intervals over a one-month period. Bacterial isolation was performed on isolated *vdac2^-/-^* individuals, healthy *vdac2^-/-^* individuals, and wild-type fish. After surface disinfection in a biosafety cabinet, dissection was performed. Organs were gently crushed using a disposable spreader to avoid contamination from digestive tract rupture. Homogenized samples were streaked onto BHI agar plates, which were then incubated upside down at 37°C for 16 hours before checking for colony growth.

A total of 12 behaviorally or morphologically abnormal *vdac2^-/-^* zebrafish were sampled, and bacterial isolation revealed massive colony growth in 10 of them (83.33%, 10/12). A small number of colonies were isolated from 37.5% (3/8) of healthy *vdac2^-/-^* zebrafish, while only 12.5% (1/8) of wild-type zebrafish showed a small number of colonies. Notably, *zfvdac2^-/-^* zebrafish larvae develop normally and remain healthy from 2.5 weeks post-fertilization (wpf) to 3 months post-fertilization (mpf). However, individual deaths gradually occur in the population after 3 mpf, and the mortality rate increases with age. This phenomenon was observed in the F2, F3, and F4 generations of zebrafish, indicating that it is a stably heritable phenotype. Bacterial isolation was performed on randomly selected dead zebrafish, and a large number of bacteria were easily isolated from the abdominal cavity of all diseased zebrafish (S4 Fig. C). After excluding laboratory-derived viral infections, the cause of death was determined to be bacterial infection based on clinical symptoms. Considering that bacteria can be isolated from the abdominal cavity of healthy *vdac2^-/-^* zebrafish and wild-type zebrafish at a certain rate without causing disease, and no outbreak of infection was observed in the *vdac2^-/-^* zebrafish population (S4 Fig. D), we speculate that *vdac2^-/-^* zebrafish may have certain defects that render them susceptible to bacterial infection. 16S rDNA-based genomic analysis revealed a large number of opportunistic pathogens in the tissues of diseased zebrafish, among which *Aeromonas sp*, *Vibrio cholerae*, and *Plesiomonas shigelloides* accounted for the highest proportions (S4 Fig. E).

Untreated diseased zebrafish usually die within a few days, but soaking in oxytetracycline aqueous solution can alleviate disease symptoms or even restore health (S4 Fig. F). However, recurrence of the disease and subsequent death occur after several weeks, suggesting the presence of potential immune disorders. It has been reported that conditional knockout of *vdac2* in mice causes excessive apoptosis and exhaustion of thymocytes (74). Zebrafish also possess an adaptive immune system composed of T cells and B cells (78), and T cells mainly develop in the thymus. It has been reported that recombination-activating protein 1 (RAG-1) expression can be detected in the thymus of wild-type zebrafish at 2-3 wpf (indicating the initiation of V(D)J recombination), and thymic involution is observed at approximately 15 wpf (79). This time point coincides with the onset of death and detection of bacterial infection in *vdac2^-/-^*zebrafish. Whether *zfvdac2* knockout affects T cell development in zebrafish warrants further investigation.

In the knockout phenotype evaluation, we found that *zfvdac2^-/-^* zebrafish had a high mortality rate within the first 7 dpf, and the dead embryos exhibited characteristics of delayed development and reduced yolk sac. VDAC2 has been widely reported to play a crucial role in apoptosis during development (25, 71, 74), which may be the cause of the high mortality rate. However, we hypothesize that the reduced yolk sac may not be related to embryonic development processes, and this phenotype may indicate defects in oocytes. It has been reported that porcine VDAC2 inhibits autophagy and promotes follicular development during folliculogenesis, and *vdac2* knockout *in vitro* enhances autophagy (77). Whether *zfvdac2* knockout affects ovarian development or oogenesis in zebrafish deserves further attention.

Xu et al. infected adult zebrafish via intraperitoneal injection of homogenate filtrate from the spleen and kidney of mandarin fish infected with ISKNV, with an inoculation dose of 25 μL per fish (2×10^11^ copies/mL). However, the initial inoculation did not cause death; lethality to zebrafish was only observed after three consecutive *in vivo* passages of the virus in zebrafish (58). In this study, we propagated ISKNV in MFF-1 cells and directly used freeze-thawed cell culture for intraperitoneal injection into zebrafish at a dose of 20 μl per fish (1×10^10^ copies/mL). At this dose, the lethality rate of ISKNV-WT to zebrafish reached 100%, while this injection dose was only 5% of that used by Xu et al. We speculate that components present in the mandarin fish homogenate may interfere with viral replication, or the determination of viral genome copy number *in vivo* may not accurately reflect the number of infectious virions, leading to a lower actual inoculation titer.

Adult *vdac2^-/-^* zebrafish over 3 mpf exhibit a certain natural mortality rate, which may affect the calculation of cumulative mortality in the challenge experiment. Through estimation, the average daily mortality rate in the *vdac2* population is approximately 0.25%, and the expected natural mortality rate within the 14-day observation period is 3.54%. This introduces a certain degree of error into the experiment; however, statistical analysis incorporating this error showed no impact on the conclusions. The thin abdominal wall of zebrafish is prone to mechanical damage and wound formation after injection, which may lead to bacterial infection. To rule out bacterial infection as a cause of zebrafish death, we added two antibiotic control groups: 20 zebrafish in each group were intraperitoneally injected with ISKNV-WT or ISKNV-Δ71 (20 μL, 1×10^10^ copies/mL) without antibiotics, while another two groups were injected with the same virus preparations containing 100 IU/mL penicillin and 100 μg/mL streptomycin. The results showed no significant difference in cumulative mortality between the antibiotic-treated and non-treated groups, indicating that antibiotics are not required in zebrafish injection experiments (data not shown). These results confirm the reliability of the data obtained from the zebrafish challenge experiments in this study.

In general, we found that *zf*VDAC2 knockout did not affect the individual pathogenicity of ISKNV and ISKNV-Δ71 in zebrafish or *zfvdac2-/-* model, which was consistent with the results of ISKNV/ISKNV-Δ71 challenge experiments on mandarin fish (21). A possible explanation is that VP71 exerts two roles in *in vivo* infection: on one hand, it has a direct pro-apoptotic function; on the other hand, VP71 binds to VDAC2 and translocates into the nucleus, increasing the sensitivity of cells to MPTP-dependent mitochondrial apoptosis. These two pathways synergistically promote cell apoptosis and enhance the virulence of ISKNV-WT. In the infection scenario where both *zf*VDAC2 and VP71 are knocked out, although the sensitivity of cells to MPTP-dependent mitochondrial apoptosis is increased, the absence of the direct pro-apoptotic effect of VP71 prevents an increase in the apoptosis rate, thus having no impact on the virulence of ISKNV-Δ71. It is worth emphasizing that mandarin fish is an extremely susceptible natural fish for ISKNV: even at a very low infection dose, mandarin fish of all sizes are highly susceptible to ISKNV (80). In contrast, in the ISKNV zebrafish infection model, zebrafish only exhibited lethality at an extremely high infection dose after multiple adaptive infections (58). Therefore, the VDAC2 knockout zebrafish model may not be an ideal choice for evaluating the real interaction between VP71 and VDAC2. The accurate verification of the interaction between VP71 and VDAC2 ultimately requires a further study based on the *vdac2* knockout mandarin fish model.

## Conclusion

This study is the first to demonstrate that ISKNV-VP71 enhances cellular apoptosis by manipulating the nuclear translocation of host VDAC2, which in turn alters mitochondrial membrane permeability. The *in vitro* experiments have essentially clarified the pathogenic mechanism of VP71. By establishing a *vdac2* knockout zebrafish infection model, we evaluated the impact of different *vdac2* phenotypes on the pathogenicity of ISKNV-WT and ISKNV-Δ71 through *in vitro* infection assays. Unexpectedly, no significant differences in the pathogenicity of ISKNV-WT and ISKNV-Δ71 were observed between the *vdac2^-/-^* and wild-type zebrafish. The complexity of ISKNV pathogenicity warrants further in-depth investigation.

## Supporting information

**S1 Fig. The whole sequence data of *mf*VDAC2 by RACE-PCR cloning.**

**S2 Fig. Cloning, Expression and Antibody Preparation of *mf*VDAC2.** (A). Prokaryotic expression and purification of *mf*VDAC2; (B). Verification of the specific recognition of *mf*VDAC2 in purified ISKNV by anti-*mf*VDAC2 antibody via WB assay.

**S3 Fig. Cross-infection experiments of ISKNV-WT or ISKNV-Δ71 on normal or *zfvdac2^⁻/⁻^* zebrafish.** Panels a, b, and c denote three independent infection experiments (N=30).

**S4 Fig. Characterization of *vdac2* knockdown zebrafish.** (A), Overview of survival statistics about *vdac2^-/^*^-^zebrafish and zebrafish wild type. X-axis was survival days. Y-axis was percentage of survival. (B), Zebrafish larvae of 4 days old were observed by stereoscopic microscope. Red arrows indicated the swim bladder. (C), The symptoms of diseased *vdac2^-/-^*zebrafish. Panel (a), the arrow indicates redness and swelling in the abdomen. Panel (b) indicate liver blood loss, ovarian ulceration, and digestive tract swelling and panel (c) indicates ascites. (D), Bacterila colony isolated by the experiment of bacterial isolation in the abdominal cavity of zebrafish. (E), bacterial identification by 16s rDNA sequencing. (F), antibiotic therapy of diseased zebrafish.

**S5 Fig. Survival rate statistics of zebrafish at 24 hpf and 7 dpf.**

**S6 Fig.** Statistics of zebrafish survival rate from 7 dpf to 300 dpf. The sample size of *vdac2^⁻/⁻^* zebrafish was 358 individuals, while that of wild - type zebrafish was 210 individuals. The horizontal axis of the coordinate system represents time, and the vertical axis represents the survival rate.

**S1 Table The primer set data used in this study.**

## Acknowledgments

This work was funded by the National Natural Science Foundation of China (Grant Number 31572642); The key areas R&D Program of Guangdong Province under No. 2021B0202040002.

## References

1. Chinchar VG, Hick P, Ince IA, Jancovich JK, Marschang R, Qin Q, et al. ICTV Virus Taxonomy Profile: *Iridoviridae*. J Gen Virol. 2017;98(5):890–1. 10.1099/jgv.0.000818 PMID: 28555546

2. Eaton HE, Ring BA, Brunetti CR. The genomic diversity and phylogenetic relationship in the family *Iridoviridae*. Viruses. 2010;2(7):1458–75. 10.3390/v2071458 PMID: 21994690

3. Kurita J, Nakajima K. Megalocytiviruses. Viruses. 2012;4(4):521–38. 10.3390/v4040521 PMID: 22590684

4. Fu W, Li Y, Fu Y, Zhang W, Luo P, Sun Q, et al. The inactivated ISKNV-I vaccine confers highly effective cross-protection against epidemic RSIV-I and RSIV-II from cultured spotted sea bass *Lateolabrax maculatus*. Microbiol Spectr. 2023;11(3):e04495–22. 10.1128/spectrum.04495-22 PMID: 37222626

5. Ramirez-Paredes JG, Paley RK, Hunt W, Feist SW, Stone DM, Field TR, et al. First detection of infectious spleen and kidney necrosis virus (ISKNV) associated with massive mortalities in farmed tilapia in Africa. Transbound Emerg Dis. 2021;68(3):1550–63. 10.1111/tbed.13825 PMID: 32920975

6. Figueiredo HCP, Tavares GC, Dorella FA, Rosa JCC, Marcelino SAC, Pierezan F, et al. First report of infectious spleen and kidney necrosis virus in Nile tilapia in Brazil. Transbound Emerg Dis. 2022;69(5):3008–15. 10.1111/tbed.14217 PMID: 34223695

7. Koda SA, Subramaniam K, Pouder DB, Yanong RP, Waltzek TB. Phylogenomic characterization of red seabream iridovirus from Florida pompano *Trachinotus carolinus* maricultured in the Caribbean Sea. Arch Virol. 2019;164(4):1209–12. 10.1007/s00705-019-04155-7 PMID: 30741339

8. Zhu Z, Duan C, Li Y, Huang C, Weng S, He J, et al. Pathogenicity and histopathology of infectious spleen and kidney necrosis virus genotype II (ISKNV-II) recovering from mass mortality of farmed Asian seabass, *Lates calcarifer*, in Southern China. Aquaculture. 2021;534(15):736326. 10.1016/j.aquaculture.2020.736326

9. Chinchar VG, Essbauer S, He JG, Hyatt A, Miyazaki T, Seligy V, et al. Family Iridoviridae. In: Fauquet CM, Mayo MA, Maniloff J, Desselberger U, Ball LA, editors. Virus taxonomy: 8th report of the International Committee on the Taxonomy of Viruses. San Diego (CA): Elsevier Academic Press; 2005. p. 163–75

10. Fu Y, Li Y, Fu W, Su H, Zhang L, Huang C, et al. Scale drop disease virus associated yellowfin seabream (*Acanthopagrus latus*) ascites diseases, Zhuhai, Guangdong, Southern China: the first description. Viruses. 2021;13(8):1617. 10.3390/v13081617 PMID: 34452481

11. de Groof A, Guelen L, Deijs M, van der Wal Y, Miyata M, Ng KS, et al. A novel virus causes scale drop disease in *Lates calcarifer*. PLoS Pathog. 2015;11(8):e1005074. 10.1371/journal.ppat.1005074 PMID: 26252390

12. Wang R, Yi Y, Liu L, Lu Y, Weng S, He J, et al. ORF005L from infectious spleen and kidney necrosis virus is located in the inner mitochondrial membrane and induces apoptosis. Virus Genes. 2014;49(2):269–77. 10.1007/s11262-014-1088-2 PMID: 24862228

13. Xu X, Weng S, Lin T, Tang J, Huang L, Wang J, et al. VP23R of infectious spleen and kidney necrosis virus mediates formation of virus-mock basement membrane to provide attaching sites for lymphatic endothelial cells. J Virol. 2010;84(22):11866–75. 10.1128/JVI.00990-10 PMID: 20810728

14. Xu X, Yan M, Wang R, Lin T, Tang J, Li C, et al. VP08R from infectious spleen and kidney necrosis virus is a novel component of the virus-mock basement membrane. J Virol. 2014;88(10):5491–501. 10.1128/JVI.03776-13 PMID: 24599992

15. He J-H, Shen W, Han D, Yan M, Luo M, Deng H, et al. Molecular mechanism of the interaction between *Megalocytivirus*-induced virus-mock basement membrane (VMBM) and lymphatic endothelial cells. J Virol. 2023;97(11):e0048023. 10.1128/jvi.00480-23 PMID: 37877715

16. Wang Z-L, Xu X-P, He B-L, Weng S-P, Xiao J, Wang L, et al. Infectious spleen and kidney necrosis virus ORF48R functions as a new viral vascular endothelial growth factor. J Virol. 2008;82(9):4371–83. 10.1128/jvi.02027-07 PMID: 18305039

17. He B-L, Yuan J-M, Yang L-Y, Xie J-F, Weng S-P, Yu X-Q, et al. The viral TRAF protein (ORF111L) from infectious spleen and kidney necrosis virus interacts with TRADD and induces caspase 8-mediated apoptosis. PLoS One. 2012;7(5):e37001. 10.1371/journal.pone.0037001 PMID: 22615868

18. Yuan J-M, He B-L, Yang L-Y, Guo C-J, Weng S-P, Li SC, et al. Interaction of infectious spleen and kidney necrosis virus ORF119L with PINCH leads to dominant-negative inhibition of integrin-linked kinase and cardiovascular defects in zebrafish. J Virol. 2015;89(1):763–75. 10.1128/jvi.01955-14 PMID: 25355883

19. Dong C-F, Xiong X-P, Shuang F, Weng S-P, Zhang J, Zhang Y, et al. Global landscape of structural proteins of infectious spleen and kidney necrosis virus. J Virol. 2011;85(6):2869–77. 10.1128/jvi.01444-10 PMID: 21209107

20. Shuang F, Luo Y, Xiong X-P, Weng S, Li Y, He J, et al. Virion proteins of an RSIV-type megalocytivirus from spotted knifejaw *Oplegnathus punctatus* (SKIV-ZJ07). Virology. 2013;437(2):89–99. 10.1016/j.virol.2012.12.017 PMID: 23352451

21. Zhang H, Qi H, Weng S, He J, Dong C. Deleting ORF71L of infectious spleen and kidney necrosis virus (ISKNV) resulted in virulence attenuation in Mandarin fish. Fish Shellfish Immunol. 2022;123:335–47. 10.1016/j.fsi.2022.02.041 PMID: 35217194

22. Benz R. Permeation of hydrophilic solutes through mitochondrial outer membranes: review on mitochondrial porins. Biochim Biophys Acta. 1994;1197(2):167–96. 10.1016/0304-4157(94)90004-3 PMID: 8031826

23. Messina A, Reina S, Guarino F, De Pinto V. VDAC isoforms in mammals. Biochim Biophys Acta. 2012;1818(6):1466–76. 10.1016/j.bbamem.2011.10.005 PMID: 22020053

24. Yamamoto T, Yamada A, Watanabe M, Yoshimura Y, Yamazaki N, Yoshimura Y, et al. VDAC1, having a shorter N-terminus than VDAC2 but showing the same migration in an SDS polyacrylamide gel, is the predominant form expressed in mitochondria of various tissues. J Proteome Res. 2006;5(12):3336–44. 10.1021/pr060291w PMID: 17137335

25. Cheng EHY, Sheiko TV, Fisher JK, Craigen WJ, Korsmeyer SJ. VDAC2 inhibits BAK activation and mitochondrial apoptosis. Science. 2003;301(5632):513–7. 10.1126/science.1083995 PMID: 12881569

26. Xu X, Decker W, Sampson MJ, Craigen WJ, Colombini M. Mouse VDAC isoforms expressed in yeast: Channel properties and their roles in mitochondrial outer membrane permeability. J Membr Biol. 1999;170(2):89–102. 10.1007/s002329900540 PMID: 10430654

27. Majumder S, Slabodnick M, Pike A, Marquardt J, Fisk HA. VDAC3 regulates centriole assembly by targeting Mps1 to centrosomes. Cell Cycle. 2012;11(19):3666–78. 10.4161/cc.21927 PMID: 22935710

28. Sampson MJ, Decker WK, Beaudet AL, Ruitenbeek W, Armstrong D, Hicks MJ, et al. Immotile sperm and infertility in mice lacking mitochondrial voltage-dependent anion channel type 3. J Biol Chem. 2001;276(42):39206–12. 10.1074/jbc.M104724200 PMID: 11507092

29. Kuwana T, Mackey MR, Perkins G, Ellisman MH, Latterich M, Schneiter R, et al. Bid, Bax, and lipids cooperate to form supramolecular openings in the outer mitochondrial membrane. Cell. 2002;111(3):331–42. 10.1016/S0092-8674(02)01036-X PMID: 12419244

30. Wei MC, Zong WX, Cheng EHY, Lindsten T, Panoutsakopoulou V, Ross AJ, et al. Proapoptotic BAX and BAK: A requisite gateway to mitochondrial dysfunction and death. Science. 2001;292(5517):727–30. 10.1126/science.1059108 PMID: 11326099

31. Nechushtan A, Smith CL, Lamensdorf I, Yoon SH, Youle RJ. Bax and Bak coalesce into novel mitochondria-associated clusters during apoptosis. J Cell Biol. 2001;153(6):1265–76. 10.1083/jcb.153.6.1265 PMID: 11402069

32. Chipuk JE, Bouchier-Hayes L, Green DR. Mitochondrial outer membrane permeabilization during apoptosis: the innocent bystander scenario. Cell Death Differ. 2006;13(8):1396–402. 10.1038/sj.cdd.4401963 PMID: 16710362

33. Bernardi P, Scorrano L, Colonna R, Petronilli V, Di Lisa F. Mitochondria and cell death. Mechanistic aspects and methodological issues. Eur J Biochem. 1999;264(3):687–701. 10.1046/j.1432-1327.1999.00725.x PMID: 10491114

34. Shimizu S, Narita M, Tsujimoto Y. Bcl-2 family proteins regulate the release of apoptogenic cytochrome c by the mitochondrial channel VDAC. Nature. 1999;399(6735):483–7. 10.1038/20959 PMID: 10365962

35. Brenner C, Cadiou H, Vieira HL, Zamzami N, Marzo I, Xie Z, et al. Bcl-2 and Bax regulate the channel activity of the mitochondrial adenine nucleotide translocator. Oncogene. 2000;19(3):329–36. 10.1038/sj.onc.1203298 PMID: 10656679

36. Baines CP, Kaiser RA, Sheiko T, Craigen WJ, Molkentin JD. Voltage-dependent anion channels are dispensable for mitochondrial-dependent cell death. Nat Cell Biol. 2007;9(5):550–5. 10.1038/ncb1575 PMID: 17417626

37. Dong C, Weng S, Shi X, Xu X, Shi N, He J. Development of a mandarin fish S*iniperca chuatsi* fry cell line suitable for the study of infectious spleen and kidney necrosis virus (ISKNV). Virus Res. 2008;135(2):273–81. 10.1016/j.virusres.2008.04.004 PMID: 18485510

38. Chen J, Fu Y, Li Y, Weng S, Wang H, He J, et al. Transferrin receptor 1 (TfR1) functions as an entry receptor for scale drop disease virus to invade the host cell via clathrin-mediated endocytosis. J Virol. 2025;99(8):e0067125. 10.1128/jvi.00671-25 PMID: 40719463

39. Imajoh M, Sugiura H, Oshima S. Morphological changes contribute to apoptotic cell death and are affected by caspase-3 and caspase-6 inhibitors during red sea bream iridovirus permissive replication. Virology. 2004;322(2):220–30. 10.1016/j.virol.2004.02.006 PMID: 15110520

40. Imajoh M, Ikawa T, Oshima S. Characterization of a new fibroblast cell line from a tail fin of red sea bream, *Pagrus major*, and phylogenetic relationships of a recent RSIV isolate in Japan. Virus Res. 2007;126(1-2):45–52. 10.1016/j.virusres.2006.12.020 PMID: 17335926

41. Perry SW, Norman JP, Barbieri J, Brown EB, Gelbard HA. Mitochondrial membrane potential probes and the proton gradient: a practical usage guide. Biotechniques. 2011;50(2):98–115. 10.2144/000113610 PMID: 21486251

42. Logan A, Pell VR, Shaffer KJ, Evans C, Stanley NJ, Robb EL, et al. Assessing the Mitochondrial Membrane Potential in Cells and In Vivo using Targeted Click Chemistry and Mass Spectrometry. Cell Metab. 2016;23(2):379–85. 10.1016/j.cmet.2015.11.014 PMID: 26712463

43. Suski JM, Lebiedzinska M, Bonora M, Pinton P, Duszynski J, Wieckowski MR. Relation between mitochondrial membrane potential and ROS formation. Methods Mol Biol. 2012;810:183–205. 10.1007/978-1-61779-382-0_12 PMID: 22057568

44. Zhong Y, Tang X, Sheng X, Xing J, Zhan W. Voltage-Dependent Anion Channel Protein 2 (VDAC2) and Receptor of Activated Protein C Kinase 1 (RACK1) Act as Functional Receptors for Lymphocystis Disease Virus Infection. J Virol. 2019;93(12):e00122–19. 10.1128/jvi.00122-19 PMID: 30918079

45. Pastorino JG, Tafani M, Rothman RJ, Marcineviciute A, Hoek JB, Farber JL. Functional consequences of the sustained or transient activation by Bax of the mitochondrial permeability transition pore. J Biol Chem. 1999;274(44):31734–9. 10.1074/jbc.274.44.31734 PMID: 10531385

46. Qian T, Nieminen AL, Herman B, Lemasters JJ. Mitochondrial permeability transition in pH-dependent reperfusion injury to rat hepatocytes. Am J Physiol. 1997;273(6):C1783–92. 10.1152/ajpcell.1997.273.6.C1783 PMID: 9435481

47. Liu Y, Tran BN, Wang F, Ounjai P, Wu JL, Hew CL. Visualization of Assembly Intermediates and Budding Vacuoles of Singapore Grouper Iridovirus in Grouper Embryonic Cells. Sci Rep. 2016;6:18696. 10.1038/srep18696 PMID: 26727547

48. Saraste A, Pulkki K. Morphologic and biochemical hallmarks of apoptosis. Cardiovasc Res. 2000;45(3):528–37. 10.1016/s0008-6363(99)00384-3 PMID: 10728374

49. Sharov VG, Todor A, Khanal S, Imai M, Sabbah HN. Cyclosporine A attenuates mitochondrial permeability transition and improves mitochondrial respiratory function in cardiomyocytes isolated from dogs with heart failure. J Mol Cell Cardiol. 2007;42(1):150–8. 10.1016/j.yjmcc.2006.09.013 PMID: 17070837

50. Elustondo PA, Nichols M, Negoda A, Thirumaran A, Zakharian E, Robertson GS, et al. Mitochondrial permeability transition pore induction is linked to formation of the complex of ATPase C-subunit, polyhydroxybutyrate and inorganic polyphosphate. Cell Death Discov. 2016;2:16070. 10.1038/cddiscovery.2016.70 PMID: 27924223

51. Neginskaya MA, Solesio ME, Berezhnaya EV, Amodeo GF, Mnatsakanyan N, Jonas EA, et al. ATP synthase C-subunit-deficient mitochondria have a small cyclosporine A-sensitive channel, but lack the permeability transition pore. Cell Rep. 2019;26(1):11–17.e2. 10.1016/j.celrep.2018.12.033 PMID: 30605668

52. Szabó I, Zoratti M. The giant channel of the inner mitochondrial membrane is inhibited by cyclosporin A. J Biol Chem. 1991;266(6):3376–9. 10.1016/S0021-9258(19)67802-6 PMID: 1847371

53. Connern CP, Halestrap AP. Recruitment of mitochondrial cyclophilin to the mitochondrial inner membrane under conditions of oxidative stress that enhance the opening of a calcium-sensitive non-specific channel. Biochem J. 1994;302(2):321–4. 10.1042/bj3020321 PMID: 7522435

54. Broekemeier KM, Pfeiffer DR. Inhibition of the mitochondrial permeability transition by cyclosporin A during long-time-frame experiments: relationship between pore opening and the activity of mitochondrial phospholipases. Biochemistry. 1995;34(50):16440–9. 10.1021/bi00050a027 PMID: 8845372

55. Broekemeier KM, Dempsey ME, Pfeiffer DR. Cyclosporin A is a potent inhibitor of the inner membrane permeability transition in liver mitochondria. J Biol Chem. 1989;264(14):7826–30 10.1016/S0021-9258(18)83116-7 PMID: 2470734

56. Petronilli V, Cola C, Massari S, Colonna R, Bernardi P. Physiological effectors modify voltage sensing by the cyclosporin A-sensitive permeability transition pore of mitochondria. 10.1016/S0021-9258(20)80631-0 J Biol Chem. 1993;268(29):21939–45. PMID: 8408050

57. Li Z, Wang Y, Xue Y, Li X, Cao H, Zheng SJ. Critical role for voltage-dependent anion channel 2 in infectious bursal disease virus-induced apoptosis in host cells via interaction with VP5. J Virol. 2012;86(3):1328–38. 10.1128/jvi.06104-11 PMID: 22114330

58. Xu X, Zhang L, Weng S, Huang Z, Lu J, Lan D, et al. A zebrafish (*Danio rerio*) model of infectious spleen and kidney necrosis virus (ISKNV) infection. Virology. 2008;376(1):1–12. 10.1016/j.virol.2007.12.026 PMID: 18423508

59. Rossi A, Kontarakis Z, Gerri C, Nolte H, Hölper S, Krüger M, et al. Genetic compensation induced by deleterious mutations but not gene knockdowns. Nature. 2015;524(7564):230–3. 10.1038/nature14580 PMID: 26168398

60. Zhu P, Ma Z, Guo L, Zhang W, Zhang Q, Zhao T, et al. Short body length phenotype is compensated by the upregulation of nidogen family members in a deleterious mutation of zebrafish. J Genet Genomics. 2017;44(11):553–6. 10.1016/j.jgg.2017.09.011 PMID: 29169924

61. Ma Z, Zhu P, Shi H, Guo L, Zhang Q, Chen Y, et al. PTC-bearing mRNA elicits a genetic compensation response via Upf3a and COMPASS components. Nature. 2019;568(7751):259–263. 10.1038/s41586-019-1057-y PMID: 30944473

62. Young MJ, Bay DC, Hausner G, Court DA. The evolutionary history of mitochondrial porins. BMC Evol Biol. 2007;7:31. 10.1186/1471-2148-7-31 PMID: 17328803

63. Qi H, Yi Y, Weng S, Zou W, He J, Dong C. Differential autophagic effects triggered by five different vertebrate iridoviruses in a common, highly permissive mandarinfish fry (MFF-1) cell model. Fish Shellfish Immunol. 2016;49:407–19. 10.1016/j.fsi.2015.12.041 PMID: 26748344

64. Jia K-T, Wu Y-Y, Liu Z-Y, Mi S, Zheng Y-W, He J, et al. Mandarin fish caveolin 1 interaction with major capsid protein of infectious spleen and kidney necrosis virus and its role in early stages of infection. J Virol. 2013;87(6):3027–38. 10.1128/jvi.00552-12 PMID: 23283951

65. Wilson JD, Bigelow CE, Calkins DJ, Foster TH. Light scattering from intact cells reports oxidative-stress-induced mitochondrial swelling. Biophys J. 2005;88(4):2929–38. 10.1529/biophysj.104.054528 PMID: 15653724

66. Madesh M, Hajnóczky G. VDAC-dependent permeabilization of the outer mitochondrial membrane by superoxide induces rapid and massive cytochrome release. J Cell Biol. 2001;155(6):1003–15. 10.1083/jcb.200105057 PMID:11739410

67. Rizzuto R, Bernardi P, Pozzan T. Mitochondria as all-round players of the calcium game. J Physiol. 2000;529(1):37–47. 10.1111/j.1469-7793.2000.00037.x PMID: 11080249

68. Maldonado EN, Lemasters JJ. ATP/ADP ratio, the missed connection between mitochondria and the Warburg effect. Mitochondrion. 2014;19 Pt A:78-84. 10.1016/j.mito.2014.09.002 PMID: 25229666

69. Fujimoto I, Pan J, Takizawa T, Nakanishi Y. Virus clearance through apoptosis-dependent phagocytosis of influenza A virus-infected cells by macrophages. J Virol. 2000;74(7):3399–403. 10.1128/jvi.74.7.3399-3403.2000 PMID: 10708457

70. Naghdi S, Hajnóczky G. VDAC2-specific cellular functions and the underlying structure. Biochim Biophys Acta. 2016;1863(10):2503–14. 10.1016/j.bbamcr.2016.04.020 PMID: 27116927

71. Chin HS, Li MX, Tan IKL, Ninnis RL, Reljic B, Scicluna K, et al. VDAC2 enables BAX to mediate apoptosis and limit tumor development. Nat Commun. 2018;9(1):4976. 10.1038/s41467-018-07309-4 PMID: 30478310

72. Weeber EJ, Levy M, Sampson MJ, Anflous K, Armstrong DL, Brown SE, et al. The role of mitochondrial porins and the permeability transition pore in learning and synaptic plasticity. J Biol Chem. 2002;277(21):18891–7. 10.1074/jbc.M201649200 PMID: 11907043

73. Shimizu H, Schredelseker J, Huang J, Lu K, Naghdi S, Lu F, et al. Mitochondrial Ca^(2+)^ uptake by the voltage-dependent anion channel 2 regulates cardiac rhythmicity. Elife. 2015;4:e04801. 10.7554/eLife.04801 PMID: 25588501

74. Ren D, Kim H, Tu H-C, Westergard TD, Fisher JK, Rubens JA, et al. The VDAC2-BAK rheostat controls thymocyte survival. Sci Signal. 2009;2(85):ra48. https://www.science.org/doi/10.1126/scisignal.2000274 PMID: 19706873

75. Wallace KN, Akhter S, Smith EM, Lorent K, Pack M. Intestinal growth and differentiation in zebrafish. Mech Dev. 2005;122(2):157–73. 10.1016/j.mod.2004.10.009 PMID: 15652704

76. Rawls JF, Mahowald MA, Goodman AL, Trent CM, Gordon JI. *In vivo* imaging and genetic analysis link bacterial motility and symbiosis in the zebrafish gut. Proc Natl Acad Sci U S A. 2007;104(18):7622–7. 10.1073/pnas.0702386104 PMID: 17456593

77. Yuan J, Zhang Y, Sheng Y, Fu XZ, Cheng HH, Zhou RJ. MYBL2 guides autophagy suppressor VDAC2 in the developing ovary to inhibit autophagy through a complex of VDAC2-BECN1-BCL2L1 in mammals. Autophagy. 2015;11(7):1081–98. 10.1080/15548627.2015.1040970 PMID: 26060891

78. Langenau DM, Zon LI. The zebrafish: a new model of T-cell and thymic development. Nat Rev Immunol. 2005;5(4):307–17. 10.1038/nri1590 PMID: 15803150

79. Lam SH, Chua HL, Gong Z, Wen Z, Lam TJ, Sin YM. Morphologic transformation of the thymus in developing zebrafish. Dev Dyn. 2002;225(1):87–94. 10.1002/dvdy.10127 PMID: 12203724

80. He J, Zeng K, Weng S, Chan S. Experimental transmission, pathogenicity and physicochemical properties of infectious spleen and kidney necrosis virus (ISKNV). Aquaculture. 2002;204(1-2):11–24. 10.1016/S0044-8486(01)00639-1

